# ’What’ and ‘where’ brain-wide pathways are dominated by internal strategies

**DOI:** 10.1101/2025.03.23.644791

**Authors:** Renana Oz Rokach, Odeya Marmor, Yoav Levy, Ariel Gilad

**Affiliations:** Department of Medical Neurobiology, Faculty of Medicine, The Institute for Medical Research Israel-Canada (IMRIC), The Hebrew University of Jerusalem, Jerusalem, Israel

## Abstract

It is long thought that higher-order sensory processing is divided into two specialized cortical streams that encode in parallel either the identity of an object or its location (i.e., ‘what’ and ‘where’ streams). Here, using the mouse whisker system, we challenge this concept by demonstrating an existence of two alternating brain-wide (beyond cortex) subnetworks that are not primarily driven by external parameters, but rather by internal strategies. We combine simultaneous brain-wide neuronal recordings in mice trained to identify or locate a certain stimulus. We find that mice deploy either an active search or a passive sensation strategy during task performance. These strategies respectively drive two distinct and brain-wide subnetworks, frontal and posterior, regardless of the type of task performed. The posterior subnetwork encoded additional internal strategic parameters such as trial history, the first task of the day, and training history. A subgroup of trials that were not dominated by frontal cortex, contained meaningful task information in the posterior cortex and several thalamic areas. Integrated together, these two subnetworks may comprise normal cognitive function.

## Introduction

Our brain constantly integrates incoming sensory information in order to provide a reliable perception which leads to an appropriate action. Still today, understanding the exact mechanism underlying sensory processing is one of the main challenges in neuroscience research. One of the most influential hypothesis related to sensory processing is the Two Stream Hypothesis which posits that the visual system in primates diverges into two parallel processing streams, a ventral stream that encodes parameters related to object identity (i.e., ‘what’ stream) and a dorsal stream that encodes information related to the location of a certain object (i.e., ‘where’ stream^1,2^). It was further suggested that these processing streams are not only related to the external parameters of the stimulus, but rather to the behavioral goal of the stimulus where the dorsal stream is motor/action related (e.g., reaching and grasping the object) whereas the ventral stream is perception related^3–5^. Directly testing these hypotheses has been rather difficult using human neuroimaging techniques which are limited to the scanner or recording from one brain area in the primate brain. Despite more than 40 years of research, the existence of two distinct processing streams along with their specific function is still hotly debated^3,6–9^. In addition, the involvement of brain areas outside the cortex within different processing streams is largely unknown.

An important aspect that may strongly affect sensory processing streams is the internal movement strategy before and during sensation. For example, moving our eyes in search of a certain object may already activate the dorsal pathway regardless of the actual position of the object. In contrast, passively waiting for a stimulus to enter the visual field may activate the ventral processing stream. In other words, internal strategies, along with external parameters, may govern the selection of distinct processing streams. Here, we use the mouse whisker system to study ‘what’ and ‘where’ processing streams, which is more advantageous than the mouse visual system. Rodents can easily localize and identify objects using their whiskers ^10,11^ and anatomical studies have highlighted a dissociation of projecting neurons into a frontal and posterior pathway diverging from posterior parietal cortex to either frontal secondary motor cortex (M2) or posterior cortex (P;^12–14)^. Association areas in the mouse are located within primary sensory areas and are multimodal, integrating information from somatosensory, auditory and visual cortices^15^. Functional studies have further highlighted a difference in encoding properties of spatial and temporal properties between posterior and frontal association areas^16–21^. In addition, it was shown that the behavioral strategies during a whisker-dependent discrimination task diverges activity to either M2 for an active strategy or P in a passive strategy^14,22^. Other factor such as trial history and motivational states have an additional effect on several posterior cortical areas, highlighting their role in higher order processing^23,25,27,29,30^. Projection-specific dissociation into frontal and posterior pathways also exists in the thalamus, emphasizing that processing pathways may involve brain areas beyond cortex^24^. These results hint of the existence of two subnetworks in the mouse brain that may also expand beyond cortex, but whether these subnetworks encode external object parameters or internal strategies is unclear.

Here, we combine wide-field imaging ^22,26,28,29,31–34^ with multi-fiber photometry ^35,36^ to simultaneously measure neuronal populations from many brain areas as the same mouse performs whisker-dependent ‘what’ and ‘where’ tasks. We find the existence of frontal and posterior subnetworks that span across many cortical and subcortical brain areas. Importantly, the recruitment of either frontal or posterior subnetwork is determined by the internal behavioral strategy of the mouse rather than the external parameters of each task.

### External tasks parameters and internal behavioral strategies

To study the ‘what’ and ‘where’ processing streams in the mouse whisker system we trained the same mouse on two whisker-dependent discrimination tasks defined as ‘what’ and ‘where’ tasks (Fig. 1a). In the ‘what’ task, mice were trained to use their whiskers in order to discriminate between two textures (i.e., object identity; Two types of sandpaper, P100 vs P1200 in grit size) where one texture is a go stimulus for which the mouse needs to lick for a reward (i.e., Hit trial), whereas the other texture the mouse is required to withhold licking (i.e., correct rejection trial, CR; Fig. 1a,b; see Methods). In the ‘where’ task, the mice learn to discriminate between different positions of the same texture (i.e., object location). Importantly, the go stimulus in both tasks was identical in terms of identity and position, enabling us to compare an identical external stimulus under different behavioral contexts (i.e., either ‘what’ or ‘where’ contexts; Fig. 1a). Mice learn the tasks sequentially (first task type alternating across mice) and reached expert level in both tasks (d’>1.5; Fig. 1c). Once mice gain expertise in one task, we imaged mesoscale neuronal dynamics during expert task performance (see below for details). After mice learned both tasks, we imaged the mice as they performed both tasks in the same day (first task type alternating across days).

**Figure 1.**
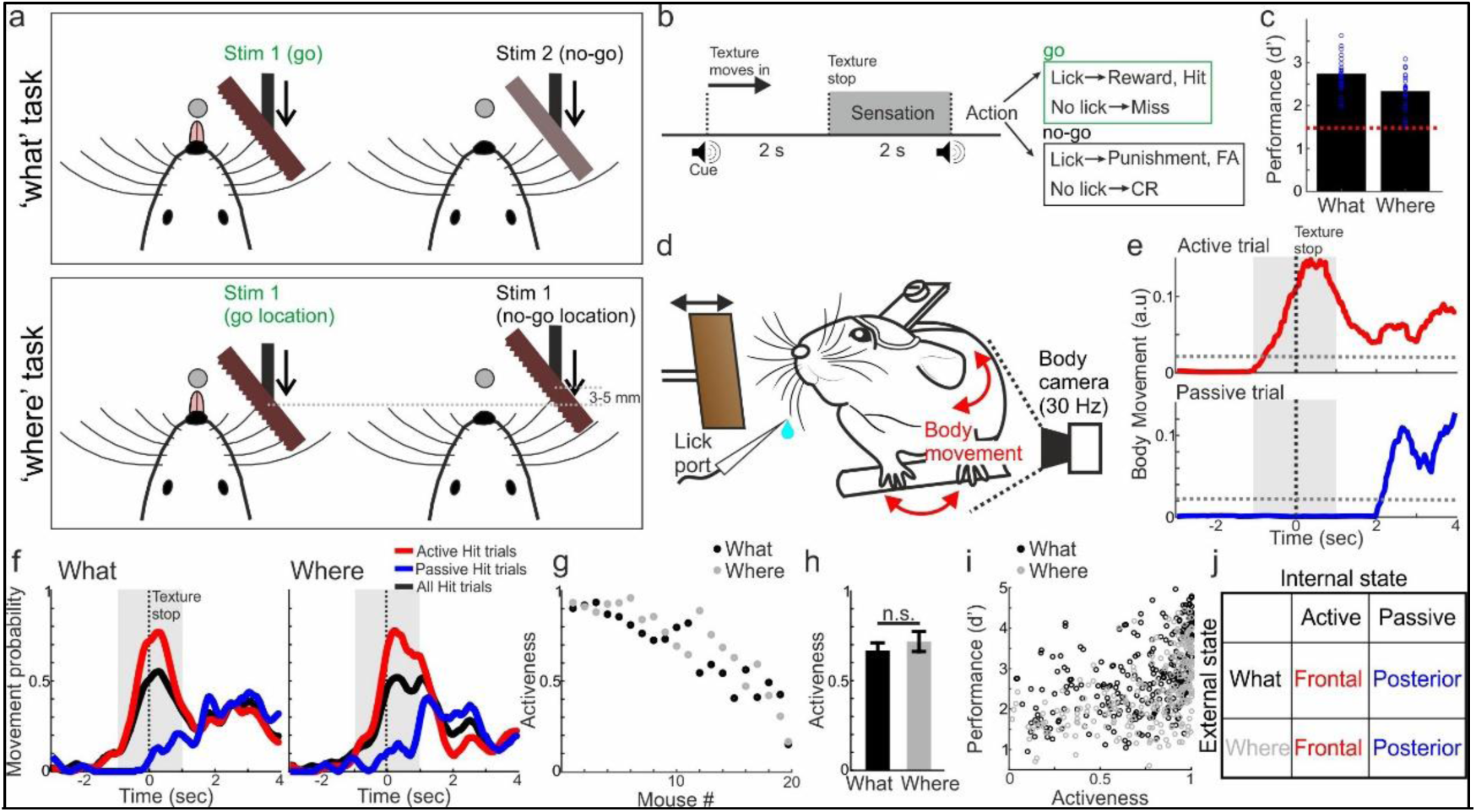
Behavioral tasks and internal strategies. **a)** Behavioral tasks. Top: ‘what’ task in which the mouse licks (go) for a specific texture. Bottom: ‘where’ task in which the mouse licks for a specific texture location. Notice that the go condition in both tasks is identical in terms of type and position. **b)** Trial structure for both tasks. **c)** behavioral performance (d’) in ‘what’ and ‘where’ tasks. Dashed red line indicates threshold for expert performance (d’=1.5). Error bars indicate mean+SEM across mice (n=19). **d)** Body movement was monitored during task performance. **e)** Example body movement of an active (top; red) and passive (bottom; blue) trial. **f)** Movement probability during an example ‘what’ (left) and ‘where’ (right) recording session for all Hit trials (black) and divided to active (red) and passive (blue) trials. **g)** Activeness for each mouse during ‘what’ (black) and ‘where’ (gray) tasks. **h)** Mean activeness during ‘what’ and ‘where’ tasks. Error bars indicate mean±SEM across mice (n=19). **i)** Performance (d’) plotted against activeness for each recording session. **j)** Overall hypothesis – frontal or posterior brain-wide subnetworks will be determined based on the internal strategy rather the task type.

During imaging experiments, we also monitored the body movements of the mouse (Fig. 1d). In general, mice deployed different movement strategies as the texture approached the whiskers. In some cases, the mouse started to actively move and whisk before the texture touched (i.e., active search strategy), whereas in other cases the mice sat quietly and waited for the texture to passively touch their whiskers (i.e., passive touch strategy). To quantify this, we divided each trial to either active (i.e., mouse moved and whisked rigorously during the sensation period; -1 to 1 second relative to texture stop) or passive (i.e., mouse quietly waited for the texture to touch the whiskers) trial (Fig. 1e;^22^). During a recording session, Hit trials can be divided into active or passive trials in a similar manner for both ‘what’ and ‘where’ tasks (Fig. 1f). We defined the activeness for each mouse as the percentage of active Hit trials during the sensation period. Activeness substantially varied across mice but was relatively similar within each mouse across both tasks, not significantly different between tasks across mice (Fig. 1g, h; p>0.05 Signed rank test). We found a significantly positive correlation between activeness and performance (r=0.47; p<0.001), although there was a substantial amount of passive recording sessions with high performance (Fig. 1i). We will later show that active and passive strategies are further linked with other non-motor internal parameters such as trial history. We hypothesize that brain-wide neuronal activity will diverge to frontal or posterior subnetworks and mainly governed by the behavioral strategy of the mouse (i.e., active or passive) and to a lesser extent the task type (i.e., either ‘what’ or ‘where’; Fig. 1j).

### Brain-wide posterior and frontal subnetworks diverge based on internal strategy

As mice performed both tasks, we obtained large-scale neuronal dynamics using different imaging preparations: 1) Wide-field imaging of the whole dorsal cortex in transgenic mice expressing a calcium indicator in layer 2/3 excitatory neurons (Fig. 2a;^22,28,37^). 2) Combined wide-field cortical imaging and multi-fiber photometry (virally injected calcium indicator) to gain simultaneous access to a wide range of brain areas within the thalamo-cortico-amygdala network (Fig. 2g). 3) Multi-fiber photometry targeting 24 cortical and subcortical recording sites including prefrontal cortex, nucleus accumbens, striatum, hippocampus, amygdala and more (Fig. 2k;^35^; see also Fig. S1). First, we focus on responses in each preparation divided into active and passive Hit trials in each task. Note that Hit trials have identical stimulus parameters (i.e., texture identity and location) in ‘what’ and ‘where’ tasks. In cortex, an active sensation map (average active during the sensation period; -1 to 1 seconds relative to texture stop) in both ‘what’ and ‘where’ tasks display enhanced activity in Barrel cortex (BC) and also in many frontal areas (Fig. 2b). In contrast, passive sensation maps display activity in BC and also posterior areas LM and P (Fig. 2b). Importantly, sensation maps are similar across ‘what’ and ‘where’ tasks for active and passive maps separately. This is emphasized when observing the difference activity map (active minus passive sensation maps), highlighting a frontal-posterior divergence based on internal strategy and reproduced across mice (Fig. 2b, c). Temporal responses display a frontal or posterior enhancement in active and passive Hit trials respectively, specifically during the sensation period (Fig. 2d). The differential response during the sensation period between active and passive trials was similar for both tasks, showing a gradient increase from negative values in posterior areas to positive values in frontal areas (Fig. 2e). It is evident that an active strategy does not only drive motor cortices but also non-motor related areas such as association areas A and AM. Grouping responses into frontal or posterior cortical areas further highlighted the differences of behavioral strategies (active or passive) but also the similarities between the what and where tasks (Fig. 2f). A two-way ANOVA of activeness (i.e., active or passive) and task type (i.e., what or where) in relation to response values (either posterior of frontal group) found an insignificant task effect, a significant activeness effect and an insignificant interaction (For posterior group – task: F(1)=2.35, p=0.13, activeness: F(1)=104.9, p<0.001, interaction: F(1)=3.11, p=0.08; For frontal group – task: F(1)=0.15, p=0.7, activeness: F(1)=2186.4, p<0.001, interaction: F(1)=0.16, p=0.69; n=6324 trials for active what, n=5733 trials for active where, n=1895 trials for passive what, n=1751 trials for passive where, taken from 6 mice). To directly compare responses between what and where tasks, we compared responses in each area within the same day and on session with a similar movement probability vector. In such cases, the differences between what and where tasks were rather minimal emphasizing that cortical responses encode task type to a lesser extent (Fig. S2). Therefore, the cortex displays a clear frontal/posterior divergence in response which is linked to behavioral strategy rather than task type.

**Figure 2.**
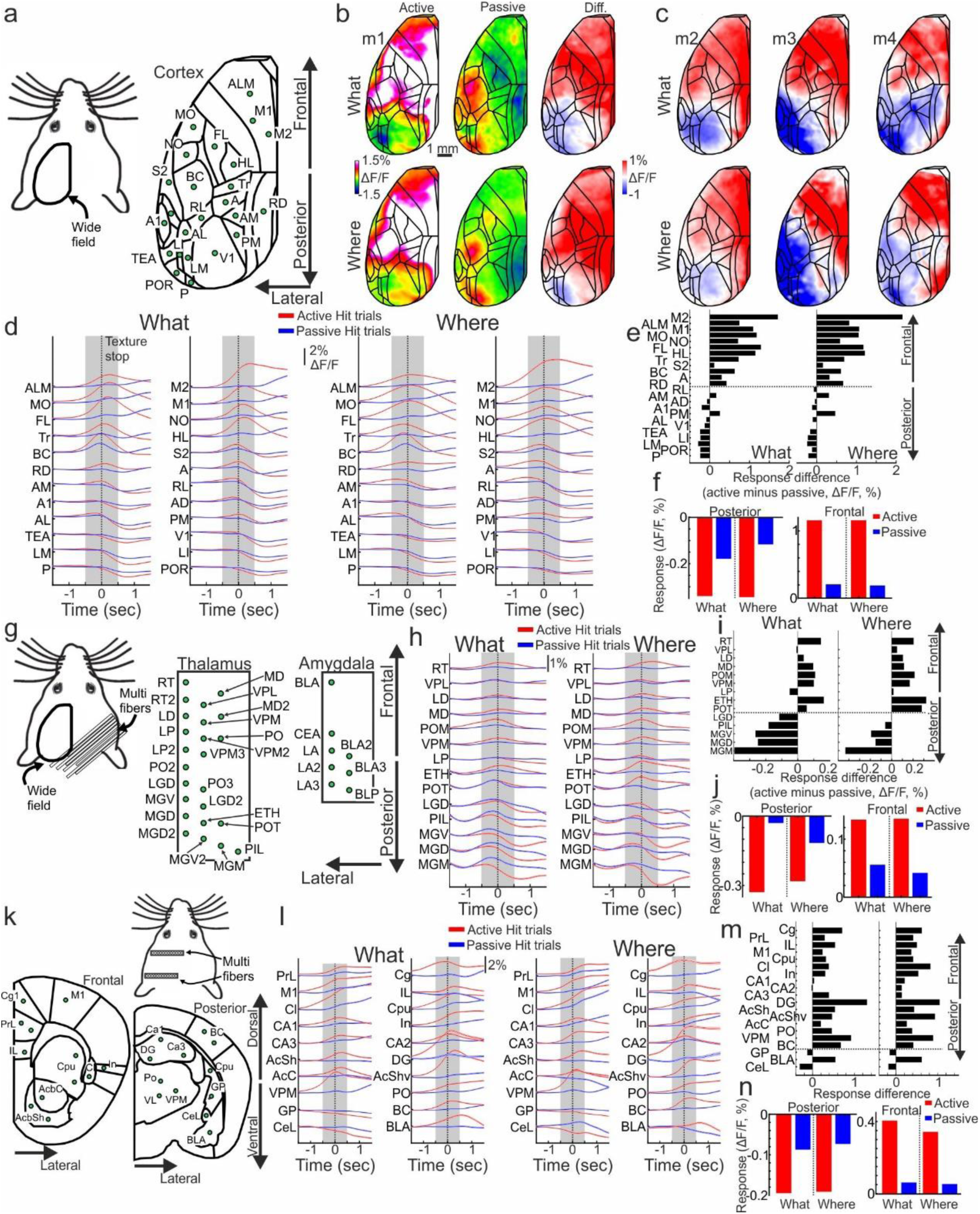
Brain-wide posterior/frontal divergence based on behavioral strategy. **a)** Wide-field imaging preparation for imaging the whole dorsal cortex. 24 Cortical brain areas were defined. **b)** Example activation maps in Hit trials (averaged within the sensation period) for active and passive states during what (top) and where (bottom) tasks. On the right is a differential activity map (active map minus passive map presented on the left). **c)** Additional example differential maps from different mice. **d)** Response in Hit trials as a function of time in each cortical area (arranged from posterior, bottom, to frontal, top) for active (red) and passive (blue) strategies in the what (left) and where (right) tasks. Shaded area indicates the sensation period. **e)** Mean differential response (active response minus passive response averaged during the sensation period) in each cortical area (arranged from posterior, bottom, to frontal, top) **f)** Mean response for active (red) and passive (blue) strategies for what and where tasks, averaged within the posterior (left) or frontal (right) brain areas (dashed line in e marks the border). **g-j)** same as d-f but for the thalamo-cortico-amygdala preparation. **k-n)** same as d-f but for the subcortical preparation. In cortex preparation: P – Posterior, POR – Post-rhinal, LM – Lateral medial, LI – Lateral intermediate, TEA – Temporal association, V1 – Primary visual, AL – Anterior lateral, PM – Posterior medial, A1 - Auditory primary, AD – Auditory dorsal, AM – Anterior medial, RL – Rostrolateral, RD – Retrosplenial dorsal, A – Anterior, BC – Barrel cortex, S2 – Secondary somatosensory, Tr, Somatosensory trunk, HL – Somatosensory hindpaw, FL – Somatosensory forepaw, NO – Somatosensory nose, MO – Somatosensory mouth, M1 – Primary motor, ALM – Anterior lateral motor, M2 – Secondary motor. In thalamo-cortico-amygdala preparation: MGM – Medial geniculate medial, MGD – Medial geniculate dorsal, MGV – Medial geniculate ventral, PIL – Posterior intralaminar, LGD - Lateral geniculate dorsal, POT - posterior thalamic, ETH – Ethmoid thalamic, LP – lateral posterior, VPM – Ventral posterior medial, POM - Posterior medial, MD – Medial dorsal, LD – Lateral dorsal, VPL – Ventral posterior lateral, RT -Reticular thalamic. BLA – Basolateral, LA – Lateral, CEA – central, BLP – Basolateral posterior. In subcortical preparation: CeL – Central amygdala, BLA – Basolateral amygdala, GP – Globus pallidus, BC – Barrel cortex, VPM – Ventral posterior medial, Po – Posterior medial, AcC – Nucleus accumbens core, AcShv – Nucleus accumbens shell ventral, AcSh – Nucleus accumbens shell, DG – Dentate gyrus, Hippocampus (cornu Ammonis) – CA1, CA2, CA3, In – Insular (In), Cl – Claustrum, Cpu – Caudate putamen, M1 – Motor primary, IL – Infralimbic, PrL – Prelimbic, Cg – Cingulate.

The thalamus follows a similar frontal-posterior divergence as the cortex. Posterior thalamic areas such as medial geniculate ventral and dorsal (MGV and MGD) display higher activity in passive trials whereas frontal thalamic areas such as reticular nucleus (RT) and posterior medial (POM) display higher activity in active trials (Fig. 2g, h). The differential response displays a gradient increase from negative values in posterior areas to positive values in frontal areas (Fig. 2h) and grouping responses into frontal or posterior parts further highlighted response differences based on strategy and to a lesser extent on task type (Fig. 2i). A two-way ANOVA of activeness (i.e., active or passive) and task type (i.e., what or where) in relation to response values (either posterior of frontal group) found a mixed task effect (significant in the posterior group but insignificant in the frontal group), a significant activeness effect and an insignificant interaction (For posterior group – task: F(1)=34.93, p<0.001, activeness: F(1)=391.02, p<0.001, interaction: F(1)=2.06, p=0.15; For frontal group – task: F(1)=1.99, p=0.16, activeness: F(1)=363.45, p<0.001, interaction: F(1)=3.03, p=0.08; n=3433 trials for active what, n=2744 trials for active where, n=2607 trials for passive what, n=2652 trials for passive where, taken from 6 mice).

In the third preparation targeting subcortical areas (Fig. 2k), we found a less pronounced frontal-posterior divergence based on activeness in which some posterior areas such as the amygdala displayed a bias towards passive trials, and many frontal areas such as prefrontal cortex and hippocampus were biased towards active trials (Fig. 2l, m). Grouping responses to posterior (amygdala and globus pallidus) and frontal groups highlighted the difference in response based on behavioral strategy but not task type (Fig. 2n). A two-way ANOVA of activeness (i.e., active or passive) and task type (i.e., what or where) in relation to response values (either posterior of frontal group) found an insignificant task effect, a significant activeness effect and an insignificant interaction (For posterior group – task: F(1)=2.83, p=0.09, activeness: F(1)=98.56, p<0.001, interaction: F(1)=2.02, p=0.16; For frontal group – task: F(1)=0.66, p=0.42, activeness: F(1)=687.26, p<0.001, interaction: F(1)=2.1, p=0.15; n=8284 trials for active what, n=8723 trials for active where, n=3467 trials for passive what, n=2948 trials for passive where, taken from 9 mice). Taken together, we find a brain-wide divergence into frontal and posterior areas that are related to active and passive behavioral strategies respectively rather than what or where tasks.

We next investigated the existence of subnetworks in the brain during task performance, with relation to either behavioral strategy or task type. In cortex, we calculated the full correlation matrix between cortical areas within the sensation period (-1 to 1 seconds relative to texture stop) separately in active and passive Hit trials (averaged across 6 mice). Next, we calculated the correlation difference matrix between active and passive trials (Fig. 3a). Positive values depict a bias towards the active strategy and negative values depict a bias towards the passive strategy. Clustering of the differential matrix revealed two distinct groups, i.e., subnetworks, that respond similarly within each group but differently between groups (Fig. 3b,c). Next, we superimposed the differential correlation values (Δr) on the cortical map for each group separately (Fig. 3d). It is evident that the two groups are divided into frontal and posterior subnetworks. Interestingly, the frontal group dominantly displays values biased towards active trials whereas the posterior group displays values that are biased towards passive trials (Fig. 3d). A two-way ANOVA of activeness (i.e., active or passive) and task type (i.e., what or where) in relation to Δr values in each group found an insignificant task effect, a significant activeness effect and an insignificant interaction (Fig. 3e; task: F(1)=0.15, p=0.7, activeness: F(1)=257.23, p<0.001, interaction: F(1)=0.39, p=0.54). A similar analysis, but subtracting ‘what’ from ‘where’ correlation matrices did not display a frontal/posterior separation (Fig. S3). These results indicate that in the cortex there are two distinct frontal and posterior subnetworks that encode active and passive strategies respectively, but not task type.

**Figure 3.**
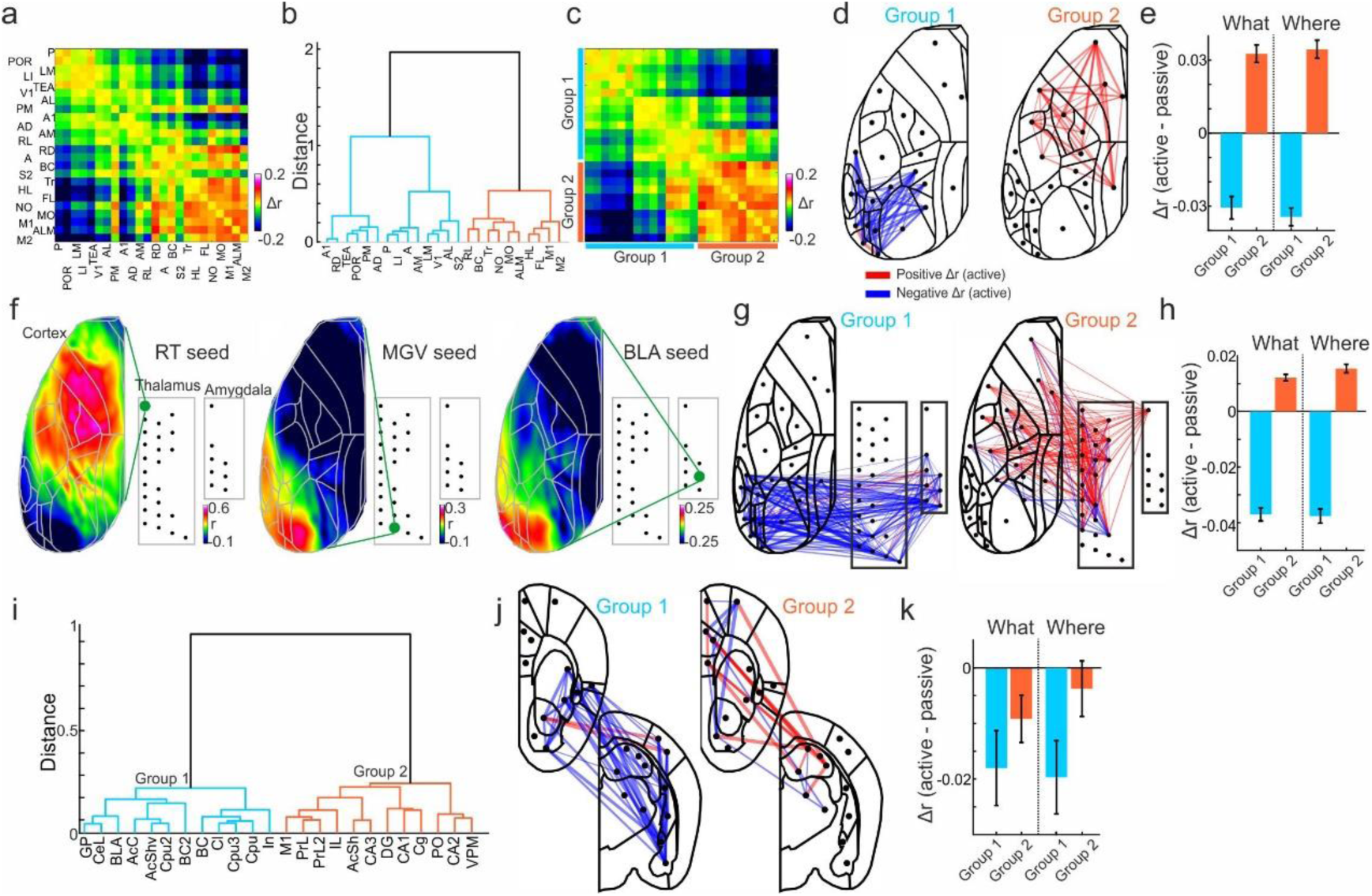
Active and passive strategies are encoded within frontal and posterior subnetworks respectively. **a-e)** In cortex. **a)** Differential correlation matrix (active correlation matrix minus passive correlation matrix) during the sensation period between all cortical areas. Data grouped together from 6 mice and both what and where tasks. Positive and negative values indicate a bias towards active and passive states respectively. **b)** Hierarchical clustering presented as a dendrogram of the differential correlation matrix in a. Two groups are highlighted in different colors. **c)** The differential correlation matrix from a sorted based on the clustering in b. d) Pairwise differential correlation values from c superimposed on the cortical atlas for each group separately. **e)** Differential correlation values averaged within group 1 (blue) or 2 (red) in the what and where tasks. Error bars depict mean±SEM across differential correlation pairs averaged across 6 mice (n=91 and 45 pairs for groups 1 and 2 respectively). **f-h)** In the thalamo-cortico-amygdala network. **f)** Seed correlation cortical map examples. Each map depicts the correlation in Hit trials during the sensation period between a seed subcortical area (either frontal RT, posterior MGV or BLA) and each pixel in the cortex. **g)** Same as d but for the thalamo-cortico-amygdala network. **h)** As in e (n=300 and 465 pairs for groups 1 and 2 respectively, averaged across 4 mice). **i-k)** In subcortex. **i)** Dendrogram as in b, but for subcortical areas. j) As in d and g. k) As in e (n=66 and 66 pairs for groups 1 and 2 respectively, averaged across 9 mice)

In the thalamo-cortico-amygdala network we find a similar strategy-dependent (rather than task-dependent) divergence into frontal and posterior subnetworks. A seed correlation map between a thalamic or amygdala area and the whole dorsal cortex (in Hit trials during the sensation period) showed that a frontal thalamic area such as RT was positively correlated with the frontal cortex (Fig. 3f). In contrast, a posterior thalamic area such as MGV or and amygdala area (BLA) were positively correlated with the posterior cortex. Across cortex, thalamus and amygdala, we find a frontal subnetwork dominated by positive Δr values (i.e., biased to an active strategy) and a posterior subnetwork displaying negative Δr values (Fig 3g). A two-way ANOVA of activeness (i.e., active or passive) and task type (i.e., what or where) in relation to Δr values in each group found an insignificant task effect, a significant activeness effect and an insignificant interaction (task: F(1)=0.59, p=0.44, activeness: F(1)=779, p<0.001, interaction: F(1)=1.19, p=0.28). These results indicate that strategy-dependent posterior/frontal divergence expands well beyond the cortex into subcortical areas such as the thalamus.

In subcortical areas, we find a similar divergence in which a posterior subnetwork, including amygdala nucleus accumbens and claustrum, displays negative Δr values (i.e., passive) whereas a frontal subnetwork, including prefrontal areas and hippocampus, displays more positive Δr values (Fig. 3j; compared to the posterior subnetwork). A two-way ANOVA of activeness (i.e., active or passive) and task type (i.e., what or where) in relation to Δr values in each group found an insignificant task effect, a significant activeness effect and an insignificant interaction (Fig. 3k; task: F(1)=0.09, p=0.76, activeness: F(1)=4.51, p=0.035, interaction: F(1)=0.37, p=0.54). Taken together, we find two brain-wide subnetworks, frontal and posterior, that are strongly linked to active and passive strategies rather that the task itself.

### Classification of choice in frontal and posterior subnetworks is determined by behavioral strategy

Next, we investigated whether these subnetworks could predict the choice of the mouse in active and passive trials. To do this, we trained a linear support vector machine (SVM) to classify between Hit and CR trials in active or passive cases and for ‘what’ or ‘where’ tasks (Fig. 4a; see Methods). This was done for each time frame separately. The accuracy of the classifier increased above chance as texture moved into touch and was higher for active compared to passive cases (Fig. 4b). On average, during the sensation period accuracy levels were similar in cortex, thalamo-cortico-amygdala network and subcortex, ranging from ∼65% to ∼75% in passive and active cases respectively (Fig. 4c). A two-way ANOVA of task and activeness in relation to accuracy found an insignificant task effect, a significant strategy effect and an insignificant interaction for cortex, thalamo-cortico-amygdala and subcortex separately (for cortex – task: F(1)=2.11, p=0.15, activeness: F(1)=330.02, p<0.001, interaction: F(1)=4, p=0.05; for thalamo-cortico-amygdala – task: F(1)=0.78, p=0.4, activeness: F(1)=38.68, p<0.001, interaction: F(1)=1.31, p=0.28; for subcortex – task: F(1)=0.54, p=0.46, activeness: F(1)=51.15, p<0.001, interaction: F(1)=2.09, p=0.15). In summary, choice can be reliably decoded in a brain-wide manner and similarly in both tasks.

**Figure 4.**
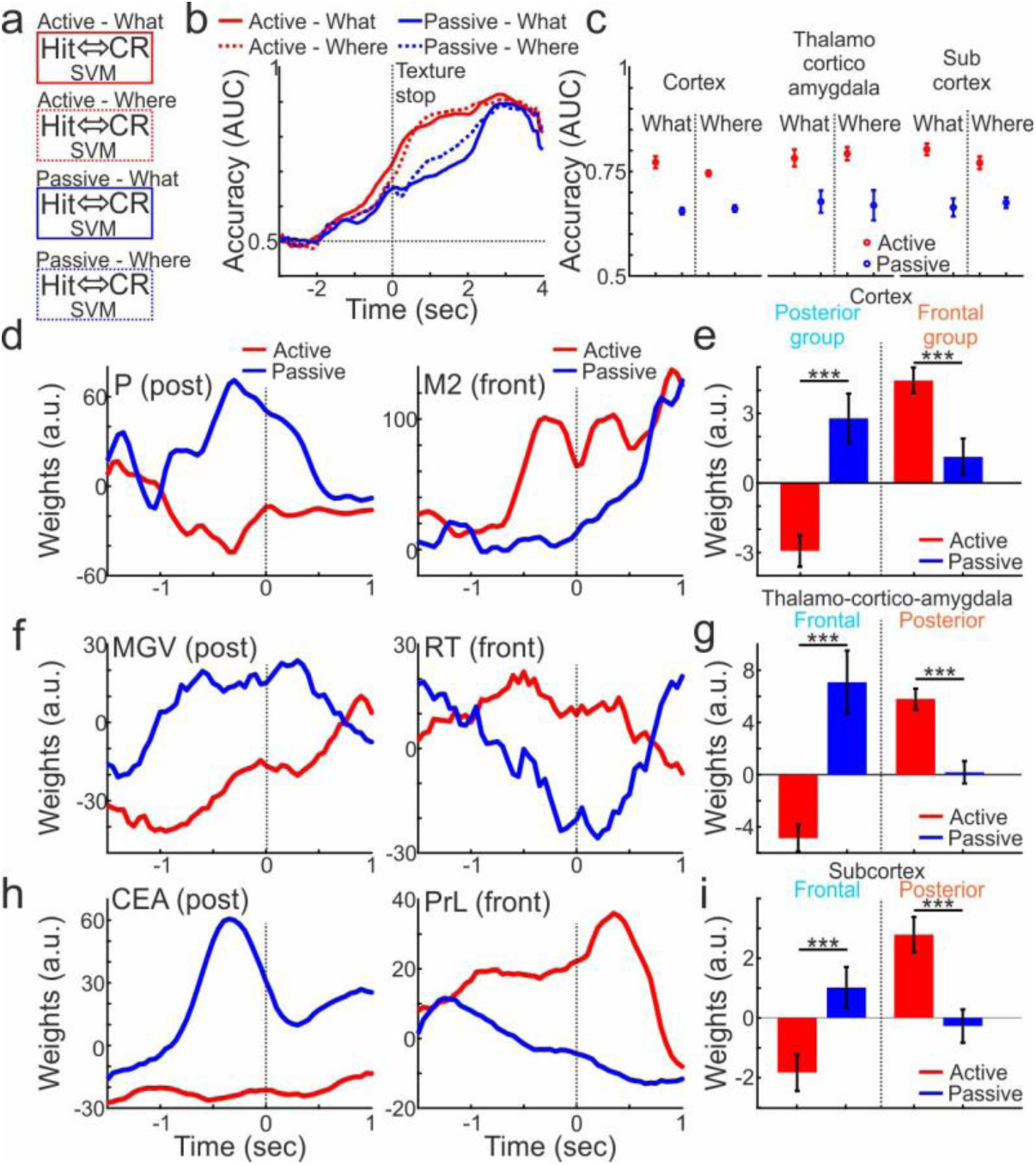
Choice decoding of active and passive strategies diverge to frontal and posterior subnetworks. **a)** A schematic of an SVM classifier decoding choice (Hit vs. CR) in different behavioral strategies (active or passive) and tasks (what or where). **b)** Classifier accuracy (averaged across 6 mice using cortical areas) as a function of time in active (red) and passive (blue) cases during what (solid) and where (dashed) tasks. **c)** Mean accuracy (averaged from -1 to 1 relative to texture stop) in active, passive, what or where using either brain areas in cortex, thalamo-cortico-amygdala, or subcortex. Error bars are SEM over mice (10 iterations from each mouse; n=6 for cortex, n=4 for thalamo-cortico-amygdala, n=9 for subcortex). **d)** Weights as a function of time during active (red) or passive (blue) strategies in an example posterior cortical area (P) and a frontal cortical area (M2). **e)** Mean weights (averaged within the sensation period) for active (red) and passive (blue) in the posterior or frontal groups for cortex. SEM are as in c. **f, g)** Similar to d and e but for the thalamo-cortico-amygdala network. **h, i)** Similar to d and e but for the subcortical network. *** - p<0.001 ranked sum test.

Next, we investigated the weights that were assigned by the classifier to each brain area or subnetwork in active and passive cases (what and where tasks were pooled together). In cortex, an example posterior area P was assigned a positive weight in the passive but not in the active case (Fig. 4d). In contrast, an example frontal area M2 was assigned positive weights in the active but not in the passive case. We next averaged the weights within the frontal or posterior subnetwork that was found in the network analysis (Fig. 3). The posterior subnetwork was assigned positive weights in the passive case which were significantly different than the negative weights in the active case (Fig. 4e; p<0.001; Rank Sum test). In the frontal subnetwork we find the opposite, significantly higher weights in the active case compared to the passive state (p<0.001; Rank Sum test).

A similar divergence of strategy-dependent frontal and posterior decoding is also present in the thalamo-cortico-amygdala network. An example posterior thalamic area (MGV) is assigned with higher weights in the passive and not the active case (Fig. 4f). In contrast, an example frontal thalamic area (RT) is assigned with higher weights in the active case and not in the passive case (Fig. 4f). The posterior subnetwork is assigned with significantly higher weights in the passive strategy compared to the active strategy (Fig. 4g; p<0.001; Rank Sum test). In contrast, the frontal subnetwork is assigned with significantly higher weights in the active strategy (p<0.001; Rank Sum test). Same is true for subcortical areas where the frontal group is assigned significantly higher weights in the active case and the posterior group is assigned significantly higher weights in the passive case (figs. 4h, i; p<0.001; Rank Sum test). This decoding divergence is similar in both ‘what’ and ‘where’ tasks separately (Fig. S4). Taken together, it is evident that choice information is encoded in frontal or posterior subnetworks depending on the internal behavioral strategy.

### Encoding of additional task-related parameters across the brain

Next, we investigated whether neuronal responses carry information related to additional behavioral and task parameters. To do this, we used a multivariate linear model to fit the neuronal response in each brain area based on a set of behavioral and task parameters (i.e., predictors) such as body movement, performance, task type, trial history and more (see Methods; Fig. 5a). The model was implemented for each brain area, time frame and mouse separately. The output of the model is a set of coefficients for each predictor in which non-zero values indicate an association between the predictor and the neuronal response in the given brain area. In cortex, we found that activeness is associated with frontal areas, but also other predictors such as performance, texture and first task (i.e., the task in which the mouse started in the day; Fig. 5b). In contrast other predictors such as history and location were more associated with posterior areas. A strong association was present specifically before and during the sensation period with some variation across predictors (Fig. 5c). A frontal/posterior divergence in coefficient values was also present to some extent within the thalamus and subcortex (Fig. 5c). Averaged across mice and divided into frontal and posterior groups activeness was significantly different between frontal and posterior groups in both cortex, thalamus and subcortex (Fig. 5d; p<0.001 Mann-Whitney U-test). In addition, we find that other parameters were significantly different between frontal and posterior, for example performance was significantly higher in frontal compared to posterior groups in both cortex, thalamus and subcortex. Surprisingly, location encoding was significantly higher in posterior groups whereas texture encoding was more related to frontal cortex. These results indicate that external parameters may be encoded within these subnetworks, but to a smaller extent. In addition, internal parameters such as the type of task in the start of the day or the history of the previous trial also displayed frontal and posterior divergence. The task predictor was also slightly significant between frontal and posterior groups, displaying mixed dynamics and complex spatiotemporal dynamics (see section below for additional analysis). The full coefficient dynamics of all predictors are presented in Fig. S5.

**Figure 5.**
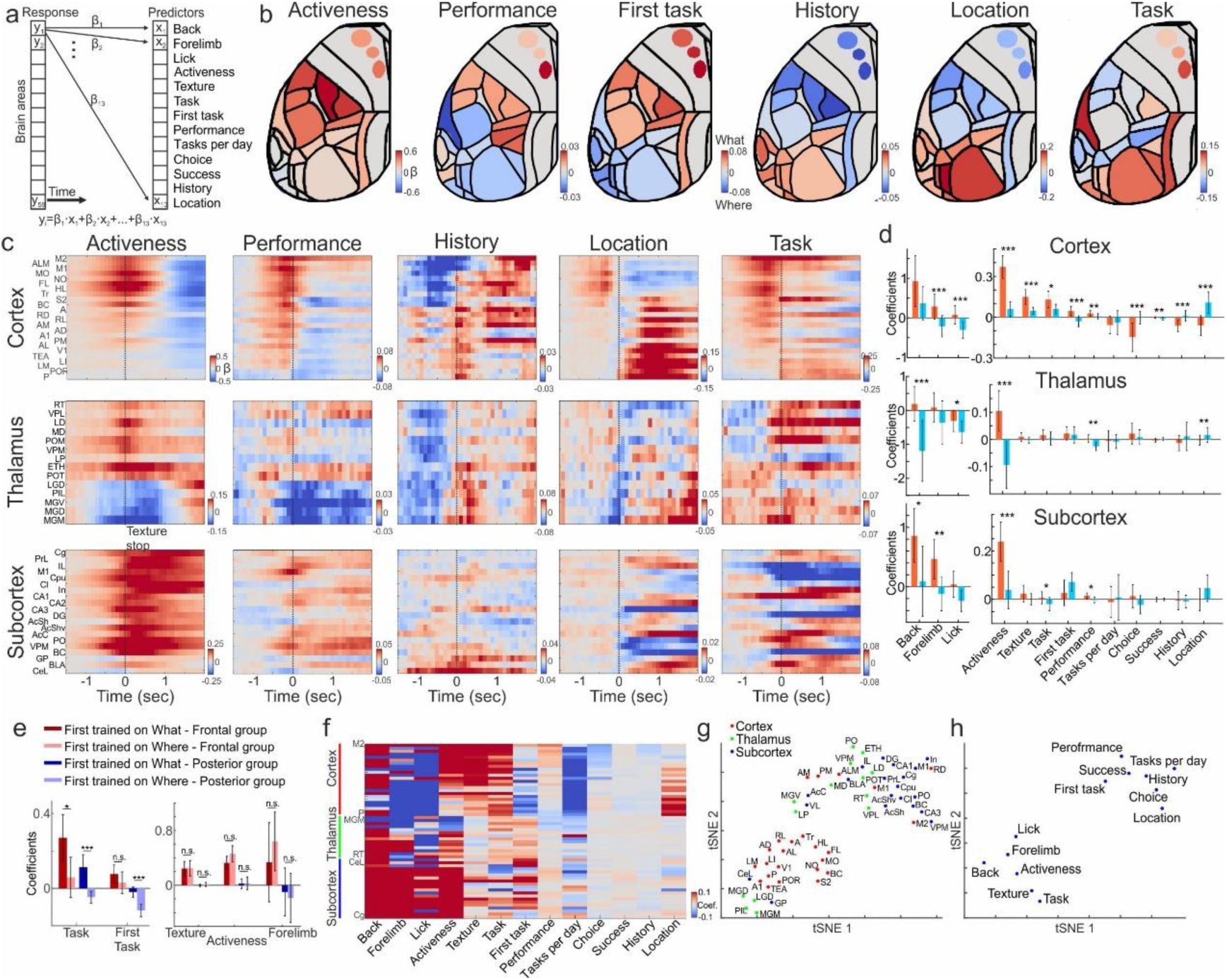
Frontal and posterior subnetworks encode additional task and behavioral parameters. **a)** A schematic of a multivariate linear model that predicts neuronal responses in each brain area (as a function of time) from a set of task and behavioral variables. **b)** Example cortical maps of coefficient values for several predictors averaged within the sensation period. Color denotes coefficient values. **c)** Temporal dynamics of example predictors (averaged across all mice) for each brain area in the cortex, thalamus or subcortex. **d)** Grand average of coefficient values for each predictor in cortex, thalamus and subcortex, grouped into frontal (red) or posterior (blue) subgroups (division as in figure 2; Averaged within the sensation period). Error bars depict S.E.M. across areas and mice (For cortex: 5 mice, 12 frontal and 12 posterior areas; For thalamus: 4 mice, 9 frontal and 5 posterior areas; For subcortex: 10 mice, 17 frontal and 3 posterior areas). **e)** Coefficients of specific predictors for cortical frontal and posterior subgroups divided into mice that either started to train on the what task (darker shades) or the where task (lighter sheds). Error bars as in d, but for 3 mice for each group. **f)** Full coefficient matrix for each brain area and predictor averaged within the sensation period. **g)** a tSNE plot of the coefficient matrix in f on the brain areas. **h)** a tSNE plot of the coefficient matrix in f on the predictors. * - p<0.05; ** - p<0.01; *** - p<0.001 ranked sum test.

Next, we divided the results based on the training history of each mice, i.e. whether the mouse started to train on the what or where task. For the mice with cortical imaging (n=6 for cortex mice, 3 started with what and 3 started with where), we find that for specific predictors, task and task first, there was a significant difference in posterior areas but not in frontal areas (p<0.001; Mann-Whitney U-test; Fig. 5e). Other predictors such as texture, activeness or forelimb, did not display a significant different in posterior or frontal groups based on the initial task the mouse was trained on (p>0.05; Mann-Whitney U-test). Taken together, these results indicate that frontal and posterior subnetworks across the brain encode additional behavioral and task related parameters and that posterior areas encode a variety of internal parameters that are related to strategic behavior.

Finally, we revisited our separation into frontal and posterior subnetworks based on the activeness of the mouse. To do this we embedded the matrix of the coefficient values for all brain areas using t-distributed stochastic neighbor embedding (t-sne; Fig. 5f). When performed on brain areas, a lower-order embedding displayed two distinct subgroups, each containing both cortical, thalamic and subcortical areas, and also strongly related to our previous division to frontal and posterior subnetworks (Fig. 5g). When performed on the predictors, a lower-order embedding displayed two distinct subgroups where one subgroup was related to activeness and movement-related predictors whereas the other group is related more internal parameters such as history and first task (Fig. 5h). Taken together, we find that frontal brain-wide subnetworks encode movement and performance related parameters whereas posterior subnetworks encode more internal and strategic parameters.

### Posterior cortical areas encode task type in a subset of trials

Up until this point, our results have indicated a frontal/posterior divergence which is governed mainly by the internal strategy of the mouse, either active or passive. In an additional effort, we wanted to investigate whether certain areas could encode the task type (either what or where) under specific circumstances. Here, we concentrated on cortex-wide data and first trained a feedforward artificial neural network (ANN) to classify trials into what or where tasks based on their spatiotemporal patterns (see Methods). We find that the model could predict task type with high accuracy for each mouse (Fig. 5a). In addition, the accuracy is significantly higher when using only posterior cortical areas compared to anterior areas (Fig. 6b; p<0.05; signed rank test; n=6 mice). These results indicate that cortical dynamics contain some sort of information regarding task type, with a bias to posterior areas.

**Figure 6.**
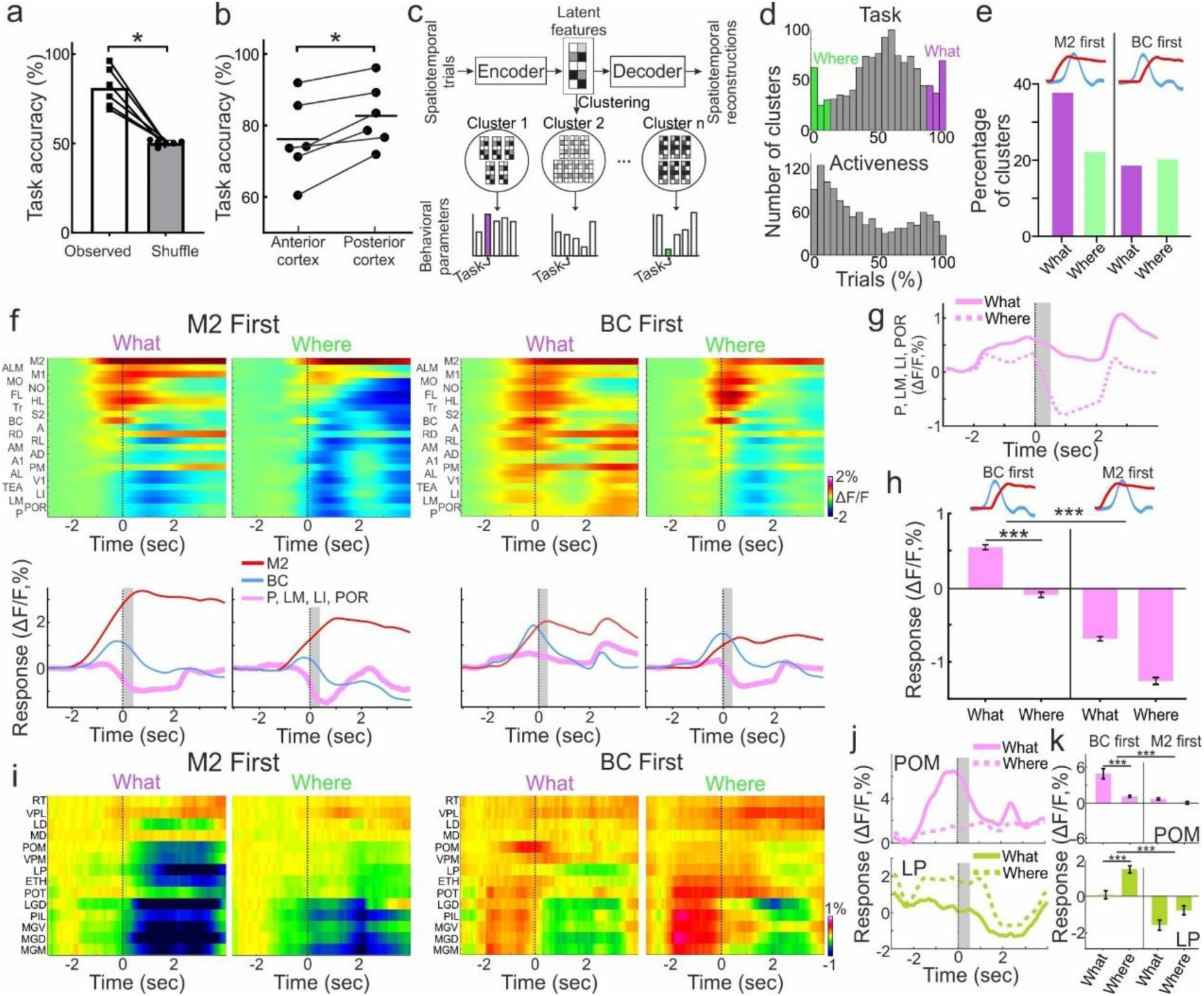
A subset of trials encodes different tasks in posterior areas. **a)** Accuracy of a feedforward ANN model trained to classify what or where tasks based on cortical dynamics. Each dot represents a mouse and is compared to trial-shuffled data. **b)** Accuracy of the model when using either only the anterior or posterior areas in the cortex. **c)** A schematic of the autoencoder model which finds reduced latent features that best represent spatiotemporal (cortical areas as a function of time) patterns. Features where clustered and specific clusters where chosen based on their behavioral parameters. **d)** Distribution for the number of clusters as a function of the percentage of trials within a cluster for behavioral variables task (task; what or where) or activeness (bottom; active or passive). We chase cluster that contained trail that where either what (purple) or where (green, more than 85% of the trials within cluster). **e)** Percentage of the chosen clusters for displaying either responses in M2 before BC or the other way around. **f)** Top: Spatiotemporal dynamics of the four groups: M2 first what, M2 first where, BC first what, BC first where. Color denotes ΔF/F. Bottom: responses in M2 (red), BC (blue) and posterior areas (P, LM, LI, POR) for each of the four groups. **g)** Reposes in posterior areas for BC first what (solid) and where (dashed) tasks. **h)** Grand average response in posterior areas for the four groups (n=1133, 1228, 2135, 832 trials for BC first what, BC first where, M2 first what, M2 first where). **i)** Same as in f, but for thalamic areas. **j)** Same as in g but for areas POM and LP in thalamus. **k)** same as in h but for areas POM and LP in thalamus. * - p<0.05; *** - p<0.001 ranked sum test.

Next, we used an autoencoder neural network model to unravel latent reduced features of spatiotemporal trials (Fig. 5c; Methods). The reduced latent features were then clustered into 1200 clusters and the profile of their behavioral parameters were assessed (Fig. 6c). The distribution of clusters is varied for each behavioral parameter, e.g., activeness displays a u-shaped distribution indicating that the model extracts a substantial number of clusters that contained either active or passive trials (Fig. 5d). In contrast, other behavioral parameters displayed a normal-like distribution indicating that the model was less successful in clustering trials based on that behavioral parameter (Fig. S6). The task parameter displayed a normal-like distribution, but there was also a substantial number of clusters at the edges of the task distribution, i.e., that contained a large proportion of mainly what or where trials. We focused on clusters that contained mainly one task type (what or where in at least 85% of trials within a cluster) and mainly Hit trial (at least 85% of trials within a cluster), resulting in 124 clusters (5328 trials) out of a total of 1200 clusters (37,275 trials, 16546 Hit trials).

Next, we further divided these clusters based on responses in specific brain areas. Based on previous results, we found that M2 activity, emerging early and related mostly to an active strategy, dominated cortical dynamics, pushing cortical responses to the frontal cortex. Therefore, we divided clusters into either M2 response preceding or following BC response, which reflects the start of the sensation period (i.e., M2 first or BC first; Fig. 6e). Our chosen clusters were distributed rather equally between the two groups, especially BC first what and where clusters (Fig. 6e). Importantly, BC first what and where clusters were similar in term of other behavioral parameters such performance, body movement and activeness. The mean spatiotemporal patterns of each group are distinctly different (Fig. 6f). M2 first groups for both what and where display a significant suppression in the posterior cortex as compared to BC first groups, especially in lateral posterior areas (LM, LI, P and POR; Fig. 6h; p<0.001 rank sum test). In BC first groups, lateral posterior areas display a significantly positive response in what group and a significantly negative drop in the where group (Fig. 6f, g, h; p<0.001 rank sum test). Other cortical areas also display some differences between what and where groups and are presented in Figure S6. For example, areas A, AM, PM and AL, which are hypothesized with where processing, showed a prominent response just after BC during BC first where group, which was less present in BC first what group. Taken together, we find that in a distinct subset of trials in which M2 did not dominate cortical responses, posterior areas could reliably encode the what task, which is partially in line with our initial hypothesis.

We performed a similar analysis on thalamic areas based on cortical responses in M2 and BC, dividing clusters into four groups as described above (42 clusters out of 1200, 631 trials). Interestingly, we find similar dynamics as in cortex in which M2 first trials displayed suppression mainly in posterior areas (Fig. 6i), whereas in BC first trials these areas were enhanced. When comparing between BC first what and where groups, we find that area POM is significantly higher in what task whereas are LP is significantly higher in the where task (Fig. 6j, k; p<0.001; Rank sum test). Within this subgroup of trials differences between what and where tasks were not organized along the frontal/posterior axis, further indicating the complex effect of internal and external parameters on brain-wide dynamics. In summary, in both cortex and thalamus we find task information in a small subset of trials which are not dominated by frontal cortex.

## Discussion

The dominant view in sensory neuroscience suggests that sensory processing is structured along distinct cortical pathways for object identity (“what”) and object location (“where”). However, our findings challenge this framework, revealing that large-scale network engagement is primarily driven by internal behavioral strategies and to a lesser extent stimulus identity or location. An active behavioral strategy engages a frontal subnetwork including cortical M2, prefrontal cortex, anterior thalamus and hippocampus whereas a passive strategy recruits a distinct posterior subnetwork including cortical areas P, posterior thalamus and amygdala (Fig. 7). Importantly, in a large subset of trials frontal/posterior divergence was similar in both ‘what’ and ‘where’ tasks, weakening the hypothesis for two separate processing streams for object identity or location^1,2^. Another previously revised interpretation for the two streams hypothesis is that the frontal subnetwork guides action whereas the posterior subnetwork encodes perception (i.e., perception-action model;^1,3,4^). Indeed, in the active case, the mouse rigorously moves its body along with activation of the frontal subnetwork, but perception, along with the presumed activation of the posterior network, is also required in order to perform the task. In the passive case, although the mouse sits rather quietly, it is expected that future licking action (e.g., planning) would still be encoded in the frontal subnetwork to some extent, since the mouse eventually performs a licking action. In addition, in previous studies we have shown that both cortical areas M2 and P are able to maintain working memory, indicating two independent networks with memory-related components^14,22^. Taken together, our data does not fully adhere to the perception-action model. Instead, we propose that sensory integration can be alternatively processed by two distinct processing streams across the brain, an active frontal stream or a passive posterior stream. The selection of the processing stream is made based on the current internal state in an alternating manner rather than in parallel.

**Figure 7.**
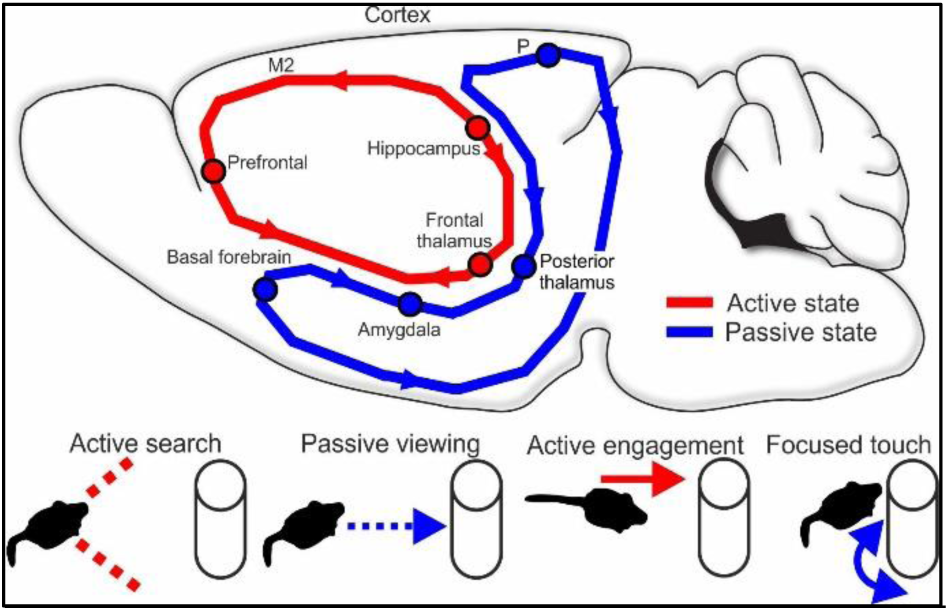
*Top:* Schematic overview of the brain-wide frontal (red) and posterior (blue) subnetworks related to active and passive states respectively. *Bottom:* Proposed natural situations in which a frontal (red) or posterior (blue) subnetwork may be implemented.

This study explores brain areas beyond the cortex, including a wide range of subcortical areas. For example, the thalamus displays an active to passive gradient along the frontal/posterior axis. A posterior subnetwork comprised of posterior cortex, posterior thalamus and amygdala is highlighted during passive sensation and may be linked to an alternative processing related to emotional circuits in the brain^38–41^. Decreased performance and prolonged learning in passive mice may further indicate that a posterior subnetwork is recruited in a more stressful context^22,28^. As for the active case, although it has been shown that movement drives brain-wide dynamics across the whole brain^42–48^, here we find a confined frontal subnetwork is recruited, whereas posterior areas often display a decrease in activity. In addition, we find that frontal and posterior subnetworks are also related to other behavioral parameters, not just activeness. The posterior subnetwork was related to internal parameters such as trial history, which task was first in the day and even which task the mouse trained on first. These relationships imply that the posterior subnetwork is linked to strategic planning and internal representations, linking the current state to previous associations based on experience.

Using an unsupervised clustering analysis, we find that a small proportion of clusters (∼10-20%) contained trials that where either what or where tasks. Within this subgroup, we find that trials in which M2 responses did not dominate (or respond rather late), posterior cortical areas indeed carry information regarding task type. Specifically, posterior areas such as LM and P displayed higher responses for what compared to where trials, in line with the hypothesis that these areas are part of the ‘what’ processing streams^52^. In addition, some thalamic higher order areas such as POM and LP also encoded task information in a small subset of trials, but with no clear frontal posterior organization. In cases in which M2 dominated cortical responses, posterior areas in cortex and thalamus were suppressed emphasizing the opposing dynamics between frontal and posterior subnetworks. We also highlight that not all trials in which M2 displayed a strong response were accompanied by strong movement, indicating that M2 is not only movement-related and may be more related to a general internal strategy. Taken together, frontal and posterior brain-wide subnetworks may carry mixed information that primarily encodes internal strategies along with some proportion of task-related what and where context.

We suggest frontal and posterior subnetworks work in a reciprocal manner in which during active sensation the frontal subnetwork takes control and has an inhibitory effect on the posterior network. A possible mechanism could be that activeness evokes a top-down control of M2 to thalamic reticular nucleus (RT), which has an inhibitory effect on many thalamic nuclei which further project to posterior cortex^49–51^. In natural conditions, the frontal subnetwork may be activated during motor execution, active search and a general unfocused state whereas the passive network can be alternatively recruited during passive viewing or focused touch, when needed to associate an object based on past experience (Fig. 7). Future studies will aim to dissect each subnetwork and investigate specific gating mechanisms and inter-subnetwork interactions underlying sensory integration in in both head-fixed and natural conditions.

## Methods

### Animals

All experiments were approved by Institutional Animal Care and Use Committee (IACUC) at the Hebrew University of Jerusalem, Israel (Permit Number: MD-20-16065-4). A total of n = 19, 8–36 week-old male mice were used in this study. In the wide-field preparation for cortex-wide imaging, we used a triple transgenic mouse line Rasgrf2-2A-dCre;CamK2a-tTA;TITL-GCaMP6f animals, expressing GCaMP6f in excitatory neocortical layer 2/3 neurons (Fig. S1;^22,28^). The rest of the mice were C57BL/6.

## Surgery

### Wide-field preparation for cortex-wide imaging

We used an intact skull preparation^53^ for chronic wide-field calcium imaging of the whole dorsal cortex^14,22,28,29^. Mice were anesthetized with 2% isoflurane (in pure O2) and body temperature was maintained at 37 °C. We applied local analgesia (lidocaine 1%), exposed and cleaned the skull, and removed some muscles to access the entire dorsal surface of the left hemisphere (∼6 × 8 mm^2^ from ∼3 mm anterior to bregma to ∼1 mm posterior to lambda; from the midline to at least 5 mm laterally). We built a wall around the hemisphere with adhesive material (iBond; UV-cured) and dental cement “worms” (Charisma). Then, we applied transparent dental cement homogenously over the imaging field (Tetric EvoFlow T1). Finally, a metal post for head fixation was glued on the back of the right hemisphere. This minimally invasive preparation enabled high-quality chronic imaging with high success rate.

### Thalamo-cortico-amygdala preparation

Here we combined wide-field cortical imaging with multi-fiber photometry to simultaneously measure calcium dynamics from the whole dorsal cortex, 24 thalamic recording sites and 8 amygdala recording sites. On the left hemisphere, we injected a calcium indicator into 13 sites across the dorsal cortex (∼150 nl of AAV2.9-hSyn-GCaMP6f) and homogeneously applied a transparent dental cement to enable wide-field imaging through the skull. On the right hemisphere, we made a small, slit-like craniotomy (from -0.82 to 3.57 mm posterior to bregma; from 1.5 to 2.63 lateral to midline) and injected the same virus into 24 thalamic sites and 8 amygdala sites with a 55-degree angle to the other hemisphere (depth ranging from 6.95 to 10.1 mm across injection sites (Fig. 2g and Fig. S1 for details). Next, we utilized fiber connector technology, specifically, ferrules that can accommodate various numbers of fibers (US Conec;^35^). Optical fibers with polyimide coating (100-μm core diameter, Thorlabs) were fitted into the ferrule, inserted into place, and sealed with dental cement. Finally, a metal post for head fixation was glued on the back of the right hemisphere. Abbreviations (from front to back): In thalamus: Reticular (RT), Ventral posterior lateral (VPL), Lateral dorsal (LD), Medial dorsal (MD), Posterior medial (POM), Ventral posterior medial (VPM), Lateral posterior (LP), Ethmoid thalamic nucleus (ETH), Posterior thalamic (POT), Lateral geniculate dorsal (LGD), Posterior intralaminar (PIL), Medial geniculate ventral (MGV), dorsal (MGD) and medial (MGM). In amygdala: Basolateral (BLA), Lateral (LA), central (CEA), Basolateral posterior (BLP).

### Subcortical preparation

For simultaneous measurements of 24 cortical and subcortical recording sites we inserted two 12-fiber implants in a frontal and posterior position (Fig. 2k and Fig. S1). We first exposed the skull, made two slit-like craniotomies (frontal slit: +1.54 mm relative to bregma, from 0.916 to 3.5 mm lateral to midline; posterior slit: -1.7 mm relative to bregma, from 1.8 to 4.384 mm lateral to midline), injected the virus with the calcium indicator, inserted the fiber implant on top (20- degree angle with fiber depths ranging from 0.375 to 4.125 mm) and sealed with dental cement. Abbreviations (from front to back): cingulate (Cg), prelimbic (PrL), infralimbic (IL), motor primary (M1), Caudate putamen (Cpu), claustrum (Cl), insular (In), CA1, CA2, CA3, dentate gyrus (DG), nucleus accumbens shell (AcbSh; ventral, AcbShv) and core (AcbC), thalamus posterior medial (Po), ventral posterior medial (VPM), barrel cortex (BC), globus pallidus (GP), basolateral (BLA) and central (CeL) amygdala.

## Imaging setups

### Wide-field imaging

The wide-field imaging setup is comprised of a sensitive CMOS camera (Hamamatsu Orca Flash 4.0 v3) mounted on top of a dual objective setup^14,22,54,55^. Two objectives (Navitar;D-5095, and D-2595) are interfaced with a dichroic (510 nm; AH) filter cube (Thorlabs). Blue LED light (Thorlabs; M470L3) is collimated and guided onto the preparation. Green light emitted from the preparation passed through both objectives before reaching the camera (20 Hz frame rate). In a subset of the imaging sessions (2 out of the 6 mice) we also interleaved and additional 405 nm light that is used as a control for non-calcium related signals (by subtracting the normalized 405 signal from the normalized 473 signal;^29,42^). In general, results were similar between corrected and non-corrected signals, as previously reported^29^, and we therefore pooled results across imaging protocols. Activation maps were registered onto the Allen atlas based on skull coordinated and functional patches (©2004 Allen Institute for Brain Science. Allen Mouse Brain Atlas. Available from: http://mouse.brain-map.org/;^13,28^).

### Multi-fiber photometry

A fiber bundle cable connects the ferrule in the mouse to the optical setup (Sylex;^35^). A Coherent OBIS LX 473-nm laser is used for excitation (and OBIS LX 405-nm as a control). Cylindrical lenses are used to create appropriate illumination patterns at the object plane of the objective. First, an the beam was expanded and collimated (GBE05-A, Thorlabs). Then, a line illumination pattern that fits the fiber array was created using two cylinder lenses (LJ1703RM-A and LJ1629RM-A, Thorlabs) placed before a dichroic cube (F38-495, AHF) and the objective (No. TL4x-SAP, Thorlabs). In a subset of the imaging sessions (3 out of the 9 mice) we also interleaved and additional 405 nm light that is used as a control for non-calcium related signals, which resulted in similar results.

### body and Whisker tracking

In addition to neuronal imaging, we tracked movements of the whiskers and the body of the mouse during the task (Fig. 1d). The mouse was illuminated with a 940-nm infrared LED. Body movements were sampled at 30 Hz (720 × 480 pixels) using a CMOS camera (The imaging source; DMK 33UX273) and an additional camera monitored the whiskers of the mouse at 30 Hz. We used movements of both forelimbs and the head/neck region to assess body movements, to reliably detect large movements (see Data Analysis below). Importantly, mice trained and performed both tasks in the dark where motor parameters were collected using infrared light.

## Behavioral tasks

### ‘What task’

Mice were trained on a go/no-go task to discriminate between two different textures (grit sizes P100: rough texture; P1200: smooth texture; 3M; Fig. 1a^11,28,29,56^). We used a data acquisition interface (USB-6001; National Instruments) and custom-written LabVIEW software (National Instruments). Each trial started with an auditory cue (stimulus cue; 2 beeps at 2 kHz, 100-ms duration with 50-ms interval), signaling the approach of either two types of sandpapers to the mouse’s whiskers as ‘go’ or ‘no-go’ textures (Fig. 1d). Sandpapers were mounted onto panels attached to a stepper motor (T-NM17A04; Zaber) mounted onto a motorized linear stage (T-LSM100A; Zaber) to move textures in and out of reach of whiskers. The texture stayed in touch with the whiskers for 2 s, and then it was moved out after which an additional auditory cue (response cue; 4 beeps at 4 kHz, 50-ms duration with 25-ms interval) signaled the start of a 2-s response period. The stimulus and response cues were identical in both textures. A water reward (∼3 μL) was given to the mouse for licking for the go texture only after the response cue (‘Hit’). Punishment with white noise was given for licking for the no-go texture (‘false alarms’; FA). Reward and punishment were omitted when mice withheld licking for the no-go (‘correct-rejections’, CR) or go (‘Misses’) textures. Note that the auditory tones merely served as cues defining the temporal trial structure, but had no predictive power with respect to go or no-go condition. Licking before the response cue was allowed and did not lead to punishment or early reward. We trained 14 mice to lick for the P100 and 5 mice to lick for the P1200.

### ‘Where task’

In this task, mice were trained on a go/no-go task to discriminate between two positions of the same texture (Fig. 1a). The position and identity of the go condition was identical to the position and identity of the go condition in the ‘what’ task. The position of the no-go condition was 3-5 mm away from the whisker pad, but still well within touch of the whiskers. The trial stucture was identical as in the ‘what’ task (i.e., same cues, stimulus period and reward window).

### Training and imaging protocol

We trained the same mouse on both tasks in a sequential manner. 11 mice started training on the ‘what’ task and 8 mice started training on the ‘where’ task. Learning the ‘what’ task started with the presentation of the go stimulus and rewarding licking during the response window. Once licking is stable, the no-go stimulus is gradually introduced (10% increase every 30-50 trials) until reaching 50% probability, mostly within the 2^nd^ day of training. On the third day, mice that persistently licked for both textures recived the no-go textures with an 80% probability until they made three consecutive correct rejections. Lerning the ‘where’ task followed a simlar protocol but starting with a relative large distance between textures (7 mm) and gradually decreasing the distance (3-5 mm) as mice learn the discrimination. Once mice became expert in the first task (d’>1.5; ∼80% performance), we started daily imaging for 4-7 days. Next, mice strated training and imaging on the second task. Once mice gained expertise in both tasks, we imaged the same mouse perfroming both tasks in the same day for an additional one to two weeks (alternatively strating with either ‘what’ or ‘where’ tasks). On average learning each task ranged from 3-8 days and the whole training and imaging protocal ranged from 24-53 days per mouse. We note that most mice performed slightly better in the ‘what’ tasks, but in both tasks performance was still high and we obtained dozens of recording days with both tasks with equal performance.

### Data analysis

Data analysis was performed using Matlab software (Mathworks). Wide-field fluorescence images were sampled down to 256 × 256 pixels and pixels outside the imaging area were discarded. This resulted in a spatial resolution of ∼40 μm/pixel and was sufficient to determine cortical borders, despite further scattering of emitted light through the tissue and skull. Each pixel and each trial the ΔF/F was calculated by dividing the raw signal by the baseline signal several frames before the stimulus cue (frame 0 division). Fiber photometry signals were collected using an identical camera and underwent the same pre-processing procedure as the wide-field signals, only that fiber tips that were focused on the camera sensor were manualy selsected and annotated for each brain area. In recording sessions where a control light (405 nm) was interleaved with the 473 nm light (sampled at 20 Hz, 10 Hz per channel), we calculated the ΔF/F for each channel separately and then subtracted the 405 control signal from the 473 signal.

### Body movement analysis

We used a body camera to detect general movements of the mouse (30 Hz frame rate). For each imaging day, we first outlined the forelimbs and the neck areas (one area of interest for each), which were reliable areas to detect general movements^14,28,29,57,58^. Next, we calculated the body movement (1 minus frame-to-frame correlation) within these areas as a function of time for each trial. Thresholding at 3 s.d. (across time frames before stimulus cue; Fig. 1e) above baseline resulted in a binary movement vector (either ‘moving’ or ‘quiet’) for each trial. The general whisker envelope was highly correlated with forelimb and back movments as previously reported^14,22,29^. By averaging across Hit trials, we derived the movement probability as a function of time (Fig. 1f black traces). The activeness for each mouse was defined as the movement probability averaged during the sensation period (-1 to 1 relative to texture stop) calculated for each task separately (Fig. 1g, h). We further divided each trial to either active, i.e., a trial in which the mouse moved for more than 0.5 seconds during the sensation period, or passive, i.e., a trial in which the mouse moved less than 0.5 seconds dutin the sensation period (Fig. 1e, f). In practice, in a typical active trial the mouse actively engaged the incoming texture, i.e, rigorously moved its forelimbs, arched its back and whisked. In contrast, in a typical passive trial the mouse quietly waited for the texture to come in contact with its whiskers. These observations led us to believe that mice deploy distinct staretgies, either active or passive, in order to perform the different tasks.

### Brain-wide correlation subnetworks

To study brain-wide subnetworks in active and passive strategies we performed the folowing: first, we calculated the full correlation matrix between all brain areas for each active or passive trial during the sensation period (pearson correlation between all areas pairs from -1 to 1 seconds relative to texture stop). This was done for each imaging preparation separately (i.e., cortex, 24 areas; thalamo-cortico-amygdala, 56 areas; subcortex, 24 areas). Next, we calculated the differential correlation matrix by subtracting the passive correlation matrix from the active correlation matrix (Fig. 3a;^26^). Positive Δr values depict a bias toward active trials whereas negative Δr values depict a bias towards the passive trials. Next, we clustered the differential matrix into two groups using a hierarchical clustering algorithm (ward method, euclidean distance) and plotted the cluster tree as a dendrogram (Fig. 3b, i; sorted differential matrices shown in Fig. 3b). Next, we used a graph plot to superimpose the Δr values on top of the brain map (Fig. 3d,g,j). Red lines indicate a bias towards active trials whereas blue lines indicate a bias towards passive trials. Thickness indicates absolute amount of Δr. This was done for the ‘what’ and ‘where’ tasks separately. Here we chose to cluster the differential correlation matrix into two groups based on our observation in cortex of a strong dissociation in activity between frontal and posterior dynamics. We note that additional smaller subnetworks could be present within the two groups.

### SVM classification of choice

To study how well does each subnetwork decode choice we trained a support vector machine (SVM) to classify trials into Hit or CR (i.e., choice) based on brain-wide neuronal dynamics^22,59^. A linear SVM was trained on either active or passive from either ‘what’ or ‘where’ tasks, i.e., four different groups of Hit and CR trials (Fig. 4a; 80% of the trials; ridge regularization; cross validation n=10; equal size groups). The remaining 20% of the trials were used as a test set for the trained SVM model and accuracy was calculated. This was done for each time frame seaprately (Fig. 4b). The weights that were assigned to each brain areas by the classifier can also be plotted as a function of time (Fig. 4d) and grouped into the two subnetworks that were clustered in the correlation analysis (Fig. 3). Our aim was to test whether choice is decoded using the frontal subnework in active trials and posterior subnetwork in the passive trials regardles of the task type.

### Multivariate linear model for linking neuronal dynamics to multiple behavioral and task parameters

To study the relationship between multiple behavioral and task parameters to neural responses in different brain areas, we applied a multiple linear regression model with Lasso regularization^42^. The goal of this analysis was to identify which parameters (other than activeness), could well predict neuronal responses in different brain areas. We selected Lasso regularization to address multicollinearity among predictors and to minimize less informative coefficients, thus simplifying the model.

For each trial, we constructed a matrix of predictive 13 variables (i.e., predictors) representing different behavioral and task parameters. These variables included binary, categorical (converted into dummy variables), and continuous variables, as follows:

1. Task: Binary variable indicating the type of task performed in the trial - 1 for the “what” task and 0 for the “where” task.
2. Location: Binary variable indicating the location of the texture stimulus - 0 for a near location (relative to whisker pad) and 1 for a far location.
3. Texture: Binary variable indicating the texture type of the stimulus - 0 for a smooth texture (P1200) and 1 for a rough texture (P100).
4. Success: Binary variable indicating trial outcome - 1 for success (i.e., Hit and correct rejection trials) and 0 for an inappropriate response (Miss and false alarm trials).
5. Number of Tasks Performed in a Day: a binary variable in which 1 indicates if the mouse performed both tasks in one day and 0 if the mouse performed only on task in that day.
6. Task First: Binary variable where 1 indicates whether the first task of the day was a ‘what’ task and 0 indicates whether the first task of the day was a ‘where’ task.
7. Activeness: Binary variable where 1 indicates whether the mouse was active and 0 indicates whether the mouse was passive in a given trial.
8. Performance: A continuous variable representing the mouse’s performance, calculated as a running d-prime (30 trial bins)
9. Back Movement: Continuous variable representing back movement, calculated by frame-to-frame correlation of a selected back area in the body camera.
10. Forelimb Movement: Continuous variable representing forelimb movement, calculated by frame-to-frame correlation of a selected forelimb area in the body camera.
11. Lick Movement: Continuous variable representing jaw and licking movement, calculated by frame-to-frame correlation of a selected jaw area in the body camera.
12. Choice: A categorial variable depicting the choice of the mouse, converted into dummy variables for “Hit,” “Miss,” “CR,” and “FA.”
13. History: A categorial variable depicting the choice of the previous trial, converted into dummy variables for “Hit,” “Miss,” “CR,” and “FA.”

These predictors were used to explain the neuronal response in each brain area separately (24 areas in cortex, 14 in thalamus and 20 in subcortical preparation). This was done for each time frame separately. Since brain activity was normally distributed, we applied a standard linear regression model.

To do this, for each mouse, brain area, and time frame, we fitted a Lasso-regularized linear regression model using the following steps:

1. Data Preprocessing: Brain activity data was standardized (z-scored) across trials.
2. Null Model: A null model (intercept-only) was fit to compute the null deviance.
3. Full Model with Lasso: The full model included all predictive variables, and Lasso regularization was applied. The optimal regularization parameter (lambda) was chosen using 10-fold cross-validation, minimizing the mean squared error (MSE). The beta coefficients from the best-fitting model (i.e., with the lowest MSE) were extracted and presented for each area separately (Fig. 5). Partial R² Calculation: To evaluate the unique contribution of each variable, we computed partial R² values for each variable. Partial R² quantifies how much variance in brain activity is uniquely explained by each predictor, accounting for the influence of other variables in the model. This is calculated using the formula:

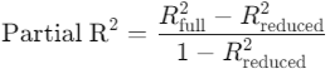

Where *R^2^_full_* is the R^2^ value from the full model, which includes all predictors. *R^2^_reduced_* is the R^2^ value from a reduced model that excludes the predictor of interest. By subtracting the R^2^ of the reduced model (which lacks the variable in question) from the R^2^ of the full model, we isolate the variance attributed to that predictor. Dividing by 1−R^2^_reduced_ adjusts for the total variance that could be explained in the absence of that predictor. Thus, partial R² gives a precise measure of each predictor’s unique explanatory power within the overall model. In general, we found that Partial R^2^ were tightly related to coefficient values, in which non-zero coefficients were associated with high Partial R^2^.

This analysis enabled us to understand how various behavioral and physiological factors influenced brain activity in different regions over time. By applying Lasso regularization, we were able to focus on the most important predictors and determine their unique contributions to brain activity.

### Task Classification Using Artificial Neural Networks (ANN)

To classify trials into task type (what or where), we trained a feedforward ANN on cortical spatiotemporal dynamics (24 cortical areas as a function of time, 70 frames for 7 seconds, for each trial) to predict whether a certain trial belongs to a what or where task. This was done for each mouse separately (n=6). The ANN was also trained separately on either only anterior or posterior cortical areas (Fig. 6b; separated as in Figure 2e). The network architecture consisted of a single hidden layer with 70 neurons, trained using cross-entropy loss minimization (patternnet function in Matlab; Epochs: 300; Early stopping: 6 validation failures; Learning rate: 0.001; L2 regularization: 0.01). The model converged using a training set (70%) and classification accuracy was calculated based on test data (30%) and cross validated (n=5). In addition to accuracy, the performance (mean squared error) of the model was 0.1886 for all brain areas, 0.2297 for anterior areas and 0.1930 for posterior areas. To ensure that classification performance was not driven by dataset-specific biases, a control analysis was performed by shuffling trial labels and re-running the classification using all brain areas.

### Autoencoder and clustering

Here, we aimed to cluster cortical spatiotemporal dynamics with an aim of finding a subgroup of trials that encodes task type (i.e., what or where). In the first step, to reduce the dimensionality an unsupervised autoencoder was trained separately for each mouse in which the input was the cortical spatiotemporal dynamics used in models above (trainAutoencoder function in MATLAB). The autoencoder architecture consisted of a single hidden layer with 300 neurons, capturing a compressed representation of the data (Fig. 6c; parameters: MaxEpochs: 200; L2 weight regularization: 0.004; Sparsity regularization: 4; Sparsity proportion: 0.1; Data scaling: Enabled). The autoencoder achieved a mean performance of (MSE) of 0.0155 (n=6 mice). The hidden layer was set to 300 neurons (∼18% of the original feature space) to ensure sufficient compression while preserving task-relevant variability in the data. This choice was further refined through an empirical tuning process, where different hidden layer sizes (ranging from 100 to 500 neurons) were tested, and 300 neurons provided the best trade-off between reconstruction accuracy and clustering performance.

Next, using hierarchical clustering the latent features extracted by the model were clustered into 1200 clusters (200 per mouse) with an average of 30.77 ± 19.31 (mean±STD) trials per cluster. This number of clusters was chosen as a compromise between unique dynamics per cluster while maintaining a rather small number of clusters. Next each cluster was labeled based on the behavioral and task parameters of its trials (same parameters as the multivariate linear model). For example, a certain cluster may contain 100 trials in which 40 of them are active, 65 are what, 35 are CR and so on. The full distribution of each parameters for all clusters are given in figures 6d and S6.

For the main analysis, we focused only on clusters that contained more than 85% of trials that were either what or where and also 80% of trials that were Hit trials (Fig. 6d colored bars). This resulted in a subgroup of 124 clusters (out of 1200) which contained 5328 trials (out of 37275 trials). Next, we divided each cluster based on its response in area M2 either rising before response in BC (M2 first) or after (BC first). This step was motivated by that during active trials M2 and other frontal areas dominated cortical dynamics, whereas in cases where M2 responses were late, cortical dynamics displayed activity in other areas. To ensure within cluster consistency in activation sequences, we confirmed that over 90% of trials within a cluster exhibited the same BC-M2 activation order. This was verified using a peak-detection approach: (1) The time of the earliest significant peaks in BC or M2 were identified for each trial. (2) The earliest rise in neural activity leading to the peak was determined using a backward search algorithm, identifying the last local minimum before a peak. (3) Trials were categorized based on whether the first detected significant activity increase occurred in BC or M2. Clusters were retained if at least 90% of trials within that cluster exhibited the same BC-M2 activation sequence only 3 clusters were excluded). This analysis resulted in four distinct groups of trials (belonging to distinct clusters): (1) What task, BC-first, (2) Where task, BC-first (3) What task, M2-first (4) Where task, M2-first (Fig. 6e for their proportion). Next, a similar analysis was done for the thalamo-cortico-amygdala dataset with a hidden layer of 700 neurons (n=3 mice), where we focused on thalamic responses in the four different groups (Fig. 6i-k).

### Statistical analysis

In general, non-parametric two-tailed statistical tests were used, Rank Sum test to compare between two medians from two populations or the Wilcoxon signed-rank test to compare a population’s median to zero (or between two paired populations). In general, measurements were independently taken from different mice and repeated across several imaging days for each mouse. Multiple group correction was used when comparing between more than two groups. Two test the effect and interaction between acitveness (active or passive) and task type (what or where) in relation to neuronal response in either frontal or posterior groups, we used a two-way ANOVA.

### Histology

Mice were given an overdose of Pental and were perfused transcardially with phosphate-buffered saline (PBS) followed by 4% paraformaldehyde (PFA) in PBS. For mice with fiber implants the skull along with the brain were post-fixed for 48 hours in 4% PFA and then extracted gently out of the skull to prevent brain rupture that can be caused by extracting multiple fibers. The extracted brains were additionally post-fixed for 12–24 hours in 4% PFA in PBS and then cryoprotected for >24 h in 30% sucrose in PBS. A subset of brains with fiber implants were scanned with magnetic resonance imaging (MRI) to better localize the fibers in the brain (7T 24 cm bore; MR Solutions, Guildford, UK). Then, 100 mm coronal slices of the entire brain were made using a vibrotome (Leica VT1000 S), mounted onto glass slides, and imaged using an epi-fluorescent microscope (Nikon SMZ25; Fig. S1). For mice with cortical imaging, we reliably detected green labeling restricted to layer 2/3 across the whole cortex, as previously reported (Fig. S1a;^22,28^). For mice with fiber implants, combining both MRI scans and brain slices, we detected fiber tips and superimposed them onto a mouse atlas (Fig. S1b-e;^60^). In cases in which not all fiber tips were identified, we estimated their location based on the other fiber tips and the measurements used to build the fiber array in advance. We report that approximately 80% of tracked fibers were correctly localized within the desired brain area.

## Acknowledgements.

We would like to thank Malak Abumadi for help with genotyping and mouse maintenance. Bar Friedmann and Shaked Yadlin for help with data analysis. Fritjof Helmchen and Philipp Bethge for help with the transgenic mouse lines. The Wohl Institute for Translational Medicine at Hadassah Hebrew University Medical Center for assistance with MRI histology. Yael Bitterman and Yasir Gallero-Salas for constructive feedback on the manuscript. This work is funded by the European Union (ERC Starting Grant, MESO-AG, 101040378) and a Hebrew University start-up grant.

## Author contributions

R.O.R and A.G. conceptualized the study. R.O.R., O.M., Y.L, and A.G. performed the experiments and preprocessed the data. R.O.R., and A.G. analyzed the data. R.O.R., and A.G. wrote the manuscript. All authors read and approved the final version of the manuscript

## Data availability

Data will be shared upon reasonable request by the corresponding author. No new material was generated in this work.

## Code availability

Custom codes for data analysis were written in MATLAB and are available from the corresponding author upon request.

## Supplementary Materials

Figs. S1 to S5

## References

1. Goodale, M.A., and Milner, A.D. (1992). Separate visual pathways for perception and action, 10.1016/0166-2236(92)90344-8 https://doi.org/10.1016/0166-2236(92)90344-8.

2. Mishkin, M., Ungerleider, L.G., and Macko, K.A. (1983). Object vision and spatial vision: two cortical pathways. Trends Neurosci. 6, 414–417. 10.1016/0166-2236(83)90190-X.

3. Goodale, M.A., and Milner, A.D. (2018). Two visual pathways – Where have they taken us and where will they lead in future? Cortex 98, 283–292. 10.1016/J.CORTEX.2017.12.002.

4. Milner, A.D., and Goodale, M.A. (2008). Two visual systems re-viewed. Neuropsychologia 46, 774–785. 10.1016/j.neuropsychologia.2007.10.005.

5. McIntosh, R.D., and Schenk, T. (2009). Two visual streams for perception and action: Current trends. Neuropsychologia 47, 1391–1396. 10.1016/j.neuropsychologia.2009.02.009.

6. Schenk, T., and McIntosh, R.D. (2010). Do we have independent visual streams for perception and action? Cogn. Neurosci. 1, 52–62. 10.1080/17588920903388950.

7. Cloutman, L.L. (2013). Interaction between dorsal and ventral processing streams: Where, when and how? Brain Lang. 127, 251–263. 10.1016/j.bandl.2012.08.003.

8. van Polanen, V., and Davare, M. (2015). Interactions between dorsal and ventral streams for controlling skilled grasp. Neuropsychologia 79, 186–191. 10.1016/j.neuropsychologia.2015.07.010.

9. Ray, D., Hajare, N., Roy, D., and Banerjee, A. (2019). Large-scale functional integration, rather than functional dissociation along dorsal and ventral streams, underlies visual perception and action. J. Cogn. Neurosci. 32, 847–861. 10.1162/jocn_a_01527.

10. Diamond, M.E., von Heimendahl, M., Knutsen, P.M., Kleinfeld, D., and Ahissar, E. (2008). “Where” and “what” in the whisker sensorimotor system. Nat. Rev. Neurosci. 9, 601–612. 10.1038/nrn2411.

11. Chen, J.L., Carta, S., Soldado-Magraner, J., Schneider, B.L., and Helmchen, F. (2013). Behaviour-dependent recruitment of long-range projection neurons in somatosensory cortex. Nature 499, 336–340. 10.1038/nature12236.

12. Wang, Q., Sporns, O., and Burkhalter, A. (2012). Network Analysis of Corticocortical Connections Reveals Ventral and Dorsal Processing Streams in Mouse Visual Cortex. J. Neurosci. 32, 4386–4399. 10.1523/JNEUROSCI.6063-11.2012.

13. Oh, S.W., Harris, J.A., Ng, L., Winslow, B., Cain, N., Mihalas, S., Wang, Q., Lau, C., Kuan, L., Henry, A.M., et al. (2014). A mesoscale connectome of the mouse brain. Nature 508, 207–214. 10.1038/nature13186.

14. Gallero-Salas, Y., Han, S., Sych, Y., Voigt, F.F., Laurenczy, B., Gilad, A., and Helmchen, F. (2021). Sensory and Behavioral Components of Neocortical Signal Flow in Discrimination Tasks with Short-Term Memory. Neuron 109, 135–148.e6. 10.1016/J.NEURON.2020.10.017.

15. Lyamzin, D., and Benucci, A. (2019). The mouse posterior parietal cortex: Anatomy and functions. Neurosci. Res. 140, 14–22. 10.1016/j.neures.2018.10.008.

16. Glickfeld, L.L., Andermann, M.L., Bonin, V., and Reid, R.C. (2013). Cortico-cortical projections in mouse visual cortex are functionally target specific. Nat. Neurosci. 16, 219–226. 10.1038/nn.3300.

17. Orlandi, J.G., Abdolrahmani, M., Aoki, R., Lyamzin, D.R., and Benucci, A. (2023). Distributed context-dependent choice information in mouse posterior cortex. Nat. Commun. 14, 1–16. 10.1038/s41467-023-35824-6.

18. Andermann, M.L., Kerlin, A.M., Roumis, D.K., Glickfeld, L.L., and Reid, R.C. (2011). Functional specialization of mouse higher visual cortical areas. Neuron 72, 1025–1039. 10.1016/j.neuron.2011.11.013.

19. Marshel, J.H., Garrett, M.E., Nauhaus, I., and Callaway, E.M. (2011). Functional specialization of seven mouse visual cortical areas. Neuron 72, 1040–1054. 10.1016/j.neuron.2011.12.004.

20. Makino, H., Ren, C., Liu, H., Kim, A.N., Kondapaneni, N., Liu, X., Kuzum, D., and Komiyama, T. (2017). Transformation of Cortex-wide Emergent Properties during Motor Learning. Neuron 94, 880–890.e8. 10.1016/j.neuron.2017.04.015.

21. Esmaeili, V., Tamura, K., Muscinelli, S.P., Modirshanechi, A., Boscaglia, M., Lee, A.B., Oryshchuk, A., Foustoukos, G., Liu, Y., Crochet, S., et al. (2021). Rapid suppression and sustained activation of distinct cortical regions for a delayed sensory-triggered motor response. Neuron 109, 2183–2201.e9. 10.1016/J.NEURON.2021.05.005.

22. Gilad, A., Gallero-salas, Y., Groos, D., Helmchen, F., Gilad, A., Gallero-salas, Y., Groos, D., and Helmchen, F. (2018). Behavioral Strategy Determines Frontal or Posterior Location of Short-Term Memory in Neocortex. Neuron 99, 814–828.e7. 10.1016/j.neuron.2018.07.029.

23. Akrami, A., Kopec, C.D., Diamond, M.E., and Brody, C.D. (2018). Posterior parietal cortex represents sensory history and mediates its effects on behaviour. Nature 554, 368–372. 10.1038/nature25510.

24. Bennett, C., Gale, S.D., Garrett, M.E., Newton, M.L., Callaway, E.M., Murphy, G.J., and Olsen, S.R. (2019). Higher-Order Thalamic Circuits Channel Parallel Streams of Visual Information in Mice. Neuron 102, 477–492.e5. 10.1016/j.neuron.2019.02.010.

25. Hattori, R., Danskin, B., Babic, Z., Mlynaryk, N., and Komiyama, T. (2019). Area-Specificity and Plasticity of History-Dependent Value Coding During Learning. Cell 177, 1858–1872.e15. 10.1016/j.cell.2019.04.027.

26. Pinto, L., Rajan, K., DePasquale, B., Thiberge, S.Y., Tank, D.W., and Brody, C.D. (2019). Task-Dependent Changes in the Large-Scale Dynamics and Necessity of Cortical Regions. Neuron, 1–15. 10.1016/j.neuron.2019.08.025.

27. Hwang, E.J., Dahlen, J.E., Mukundan, M., and Komiyama, T. (2017). History-based action selection bias in posterior parietal cortex. Nat. Commun. 8, 1–14. 10.1038/s41467-017-01356-z.

28. Gilad, A., and Helmchen, F. (2020). Spatiotemporal refinement of signal flow through association cortex during learning. Nat. Commun. 11, 1–14. 10.1038/s41467-020-15534-z.

29. Marmor, O., Pollak, Y., Doron, C., Helmchen, F., and Gilad, A. (2023). History information emerges in the cortex during learning. Elife 12. 10.7554/ELIFE.83702.

30. Matteucci, G., Guyoton, M., Mayrhofer, J.M., Auffret, M., Foustoukos, G., Petersen, C.C.H., and El-Boustani, S. (2022). Cortical sensory processing across motivational states during goal-directed behavior. Neuron 110, 4176–4193.e10. 10.1016/j.neuron.2022.09.032.

31. Mohajerani, M.H., Chan, A.W., Mohsenvand, M., LeDue, J., Liu, R., McVea, D.A., Boyd, J.D., Wang, Y.T., Reimers, M., and Murphy, T.H. (2013). Spontaneous cortical activity alternates between motifs defined by regional axonal projections. Nat. Neurosci. 16, 1426–1435. 10.1038/nn.3499.

32. Clancy, K.B., Orsolic, I., and Mrsic-Flogel, T.D. (2019). Locomotion-dependent remapping of distributed cortical networks. Nat. Neurosci. 22, 778– 786. 10.1038/s41593-019-0357-8.

33. Allen, W.E., Kauvar, I. V., Chen, M.Z., Richman, E.B., Yang, S.J., Chan, K., Gradinaru, V., Deverman, B.E., Luo, L., and Deisseroth, K. (2017). Global Representations of Goal-Directed Behavior in Distinct Cell Types of Mouse Neocortex. Neuron 94, 891–907.e6. 10.1016/j.neuron.2017.04.017.

34. Ferezou, I., Haiss, F., Gentet, L.J., Aronoff, R., Weber, B., and Petersen, C.C.H. (2007). Spatiotemporal Dynamics of Cortical Sensorimotor Integration in Behaving Mice. Neuron 56, 907–923. 10.1016/j.neuron.2007.10.007.

35. Sych, Y., Chernysheva, M., Sumanovski, L.T., and Helmchen, F. (2019). High-density multi-fiber photometry for studying large-scale brain circuit dynamics. Nat. Methods 16, 553–560. 10.1038/s41592-019-0400-4.

36. Sych, Y., Fomins, A., Novelli, L., and Helmchen, F. (2022). Dynamic reorganization of the cortico-basal ganglia-thalamo-cortical network during task learning. Cell Rep. 40, 111394. 10.1016/j.celrep.2022.111394.

37. Madisen, L., Zwingman, T.A., Sunkin, S.M., Oh, S.W., Zariwala, H.A., Gu, H., Ng, L.L., Palmiter, R.D., Hawrylycz, M.J., Jones, A.R., et al. (2010). A robust and high-throughput Cre reporting and characterization system for the whole mouse brain. Nat. Neurosci. 13, 133–140. 10.1038/nn.2467.

38. Taylor, J.A., Hasegawa, M., Benoit, C.M., Freire, J.A., Theodore, M., Ganea, D.A., Innocenti, S.M., Lu, T., and Gründemann, J. (2021). Single cell plasticity and population coding stability in auditory thalamus upon associative learning. Nat. Commun. 12, 1–14. 10.1038/s41467-021-22421-8.

39. LeDoux, J.E., Sakaguchi, A., and Reis, D.J. (1983). Subcortical Conditioned Projections of the Medial Nucleus Mediate Emotional Responses. J. Neurosci. 4, 683–698.

40. Iwata, J., LeDoux, J.E., Meeley, M.P., Arneric, S., and Reis, D.J. (1986). Intrinsic neurons in the amygdaloid field projected to by the medial geniculate body mediate emotional responses conditioned to acoustic stimuli. Brain Res. 383, 195–214. 10.1016/0006-8993(86)90020-X.

41. Khalil, V., Faress, I., Mermet-Joret, N., Kerwin, P., Yonehara, K., and Nabavi, S. (2023). Subcortico-amygdala pathway processes innate and learned threats. Elife 12, 1–26. 10.7554/eLife.85459.

42. Musall, S., Kaufman, M.T., Juavinett, A.L., Gluf, S., and Churchland, A.K. (2019). Single-trial neural dynamics are dominated by richly varied movements. Nat. Neurosci. 22, 1677–1686. 10.1038/s41593-019-0502-4.

43. Stringer, C., Pachitariu, M., Steinmetz, N., Reddy, C.B., Carandini, M., and Harris, K.D. (2019). Spontaneous behaviors drive multidimensional, brainwide activity. Science 364, 255. 10.1126/science.aav7893.

44. Steinmetz, N.A., Zatka-Haas, P., Carandini, M., and Harris, K.D. (2019). Distributed coding of choice, action and engagement across the mouse brain. Nature 576, 266–273. 10.1038/s41586-019-1787-x.

45. Benson, B., Benson, J., Birman, D., Bonacchi, N., Carandini, M., Catarino, J.A., Chapuis, G.A., Churchland, A.K., Dan, Y., Dayan, P., et al. (2023). A Brain-Wide Map of Neural Activity during Complex Behaviour. bioRxiv, 2023.07.04.547681.

46. Khilkevich, A., Lohse, M., Low, R., Orsolic, I., Bozic, T., Windmill, P., and Mrsic-Flogel, T.D. (2024). Brain-wide dynamics linking sensation to action during decision-making. Nature, 48–50. 10.1038/s41586-024-07908-w.

47. Chen, S., Liu, Y., Wang, Z.A., Colonell, J., Liu, L.D., Hou, H., Tien, N.W., Wang, T., Harris, T., Druckmann, S., et al. (2024). Brain-wide neural activity underlying memory-guided movement. Cell 187, 676–691.e16. 10.1016/j.cell.2023.12.035.

48. Bondy, A., Charlton, J., Luo, T., Kopec, C., Stagnaro, W., Venditto, S., Lynch, L., Janarthanan, S., Oline, S., Harris, T., et al. (2024). Coordinated cross-brain activity during accumulation of sensory evidence and decision commitment. bioRxiv, 1–44.

49. Hádinger, N., Bősz, E., Tóth, B., Vantomme, G., Lüthi, A., and Acsády, L. (2023). Region-selective control of the thalamic reticular nucleus via cortical layer 5 pyramidal cells. Nat. Neurosci. 26, 116–130. 10.1038/s41593-022-01217-z.

50. Pinault, D. (2004). The thalamic reticular nucleus: Structure, function and concept 10.1016/j.brainresrev.2004.04.008.

51. Wimmer, R.D., Schmitt, L.I., Davidson, T.J., Nakajima, M., Deisseroth, K., and Halassa, M.M. (2015). Thalamic control of sensory selection in divided attention. Nature 526, 705–709. 10.1038/nature15398.

52. Wang, Q., Gao, E., and Burkhalter, A. (2011). Gateways of ventral and dorsal streams in mouse visual cortex. J. Neurosci. 31, 1905–1918. 10.1523/JNEUROSCI.3488-10.2011.

53. Silasi, G., Xiao, D., Vanni, M.P., Chen, A.C.N., and Murphy, T.H. (2016). Intact skull chronic windows for mesoscopic wide-field imaging in awake mice. J. Neurosci. Methods 267, 141–149. 10.1016/j.jneumeth.2016.04.012.

54. Abdelfattah, A.S., Ahuja, S., Akkin, T., Rao Allu, S., Brake, J., Boas, D.A., Buckley, E.M., Campbell, R.E., Chen, A.I., Cheng, X., et al. (2022). Neurophotonic tools for microscopic measurements and manipulation: status report. 10.1117/1.NPh.9.S1.013001 9, 013001. https://doi.org/10.1117/1.NPH.9.S1.013001.

55. Gilad, A. (2024). Wide-field imaging in behaving mice as a tool to study cognitive function. Neurophotonics 11, 1–16. 10.1117/1.nph.11.3.033404.

56. Levitan, D., and Gilad, A. (2024). Amygdala and Cortex Relationships during Learning of a Sensory Discrimination Task. J. Neurosci. 44, 1–13. 10.1523/JNEUROSCI.0125-24.2024.

57. Gilad, A., Gallero-Salas, Y., Groos, D., Helmchen, F., Gilad, A., Gallero-Salas, Y., Groos, D., and Helmchen, F. (2018). Behavioral Strategy Determines Frontal or Posterior Location of Short-Term Memory in Neocortex. Neuron 99, 814–828.e7. 10.1016/j.neuron.2018.07.029.

58. Gilad, A., Maor, I., and Mizrahi, A. (2020). Learning-related population dynamics in the auditory thalamus. Elife 9, 1–18. 10.7554/eLife.56307.

59. Musall, S., Sun, X.R., Mohan, H., An, X., Gluf, S., Li, S.J., Drewes, R., Cravo, E., Lenzi, I., Yin, C., et al. (2023). Pyramidal cell types drive functionally distinct cortical activity patterns during decision-making. Nat. Neurosci. 26, 495–505. 10.1038/s41593-022-01245-9.

60. Paxinos, G., and Franklin, K.B.J. (2001). Paxinos and Franklin’s the Mouse Brain in Stereotaxic Coordinates 10.1364/OE.20.020998.

