## Supplementary figures for "’What’ and ‘where’ brain-wide pathways are dominated by internal strategies"

**Supplemental figure:**

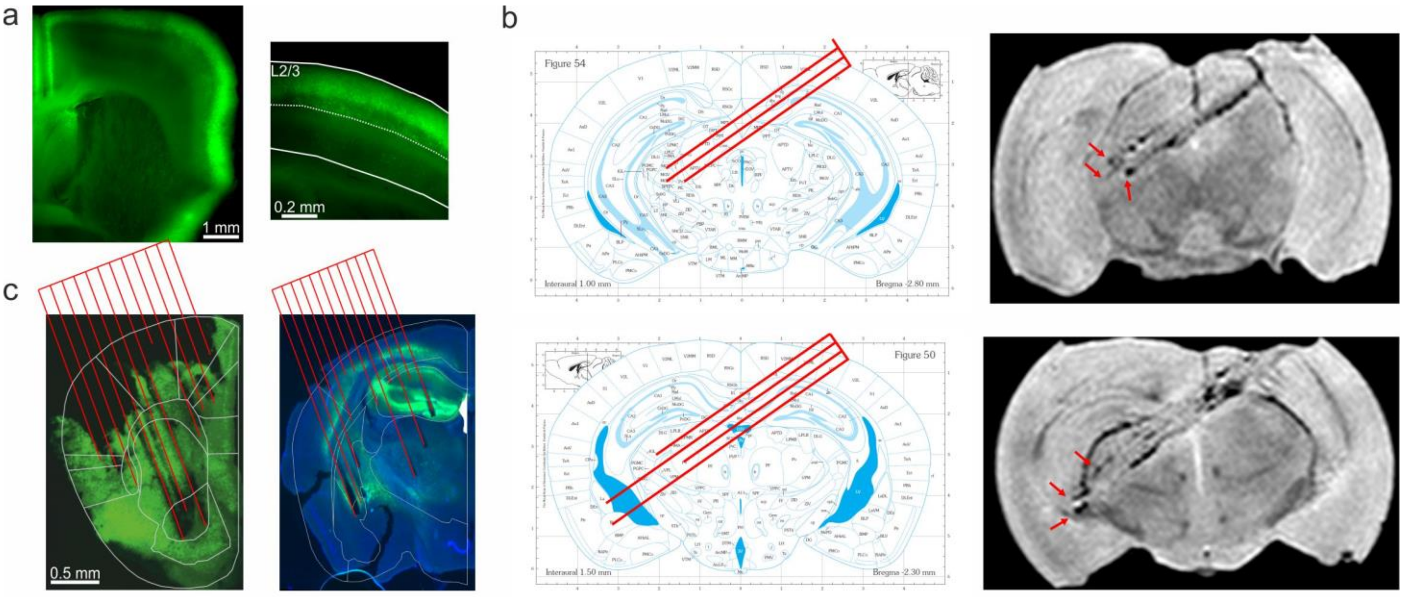


**Figure S1. Histology and fiber locations. a)** For wide-filed cortical imaging: example coronal slice displaying GCaMP6f expression specifically for layer 2/3 in cortex. These mice were used for widefield imaging in cortex. **b)** For Thalamo-cortico-amygdala preparation: *Left:* examples of planned multi-fiber array superimposed on a specific coronal slice of the Paxinos mouse atlas. *Right:* a complementary MRI scan (coronal snapshot) displaying several fiber tracks that are similar to the planned fiber coordinates on the left. Red arrows mark fiber tips. This preparation enabled to measure the thalamo-cortico-amygdala network. **c)** For subcortical preparation: coronal slices for the anterior (left; +1.54 mm relative to Bregma) and posterior (right; -1.7 mm relative to Bregma) fiber arrays used for the subcortical preparation. Fiber arrays are superimposed in red.

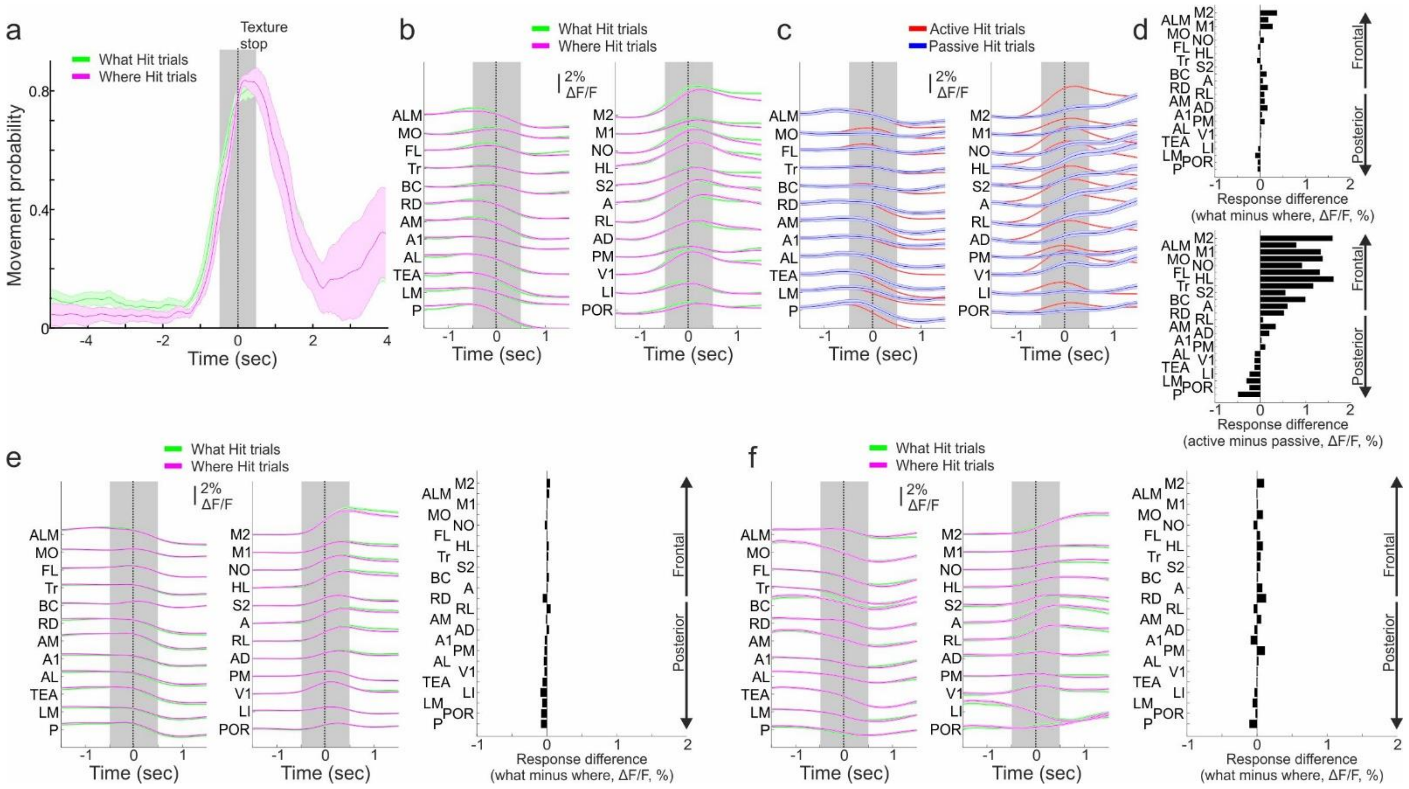


**Figure S2. Responses comparisons between 'what' and 'where' tasks. a)** Movement probability in a subset of 'what' (green) and 'where' (magenta) recording sessions taken from the same day in one mouse. Sessions were chosen based on similar movement probability across tasks. Error bars indicate mean±SEM across recording days (n=5). **b)** Response in Hit trials as a function of time in each cortical area (arranged from posterior, bottom, to frontal, top) for what (green) and where (magenta) tasks from the sessions in a. Error bars indicate mean±SEM across trials (n=496 and 375 in what and where tasks respectively). **c)** Same dataset as in b, but now divided into active (red) and passive (blue) trials (n=730 and 55 in active and passive strategies respectively). **d)** Mean differential response averaged during the sensation period (gray bar in b) for either 'what' minus 'where' responses (bottom; data in b) or active minus passive responses (bottom; data in c) in each cortical area (arranged from posterior, bottom, to frontal, top). **e,f)** same as b and d (top) but for two additional mice. In general, response during what and where tasks are similar across cortex.

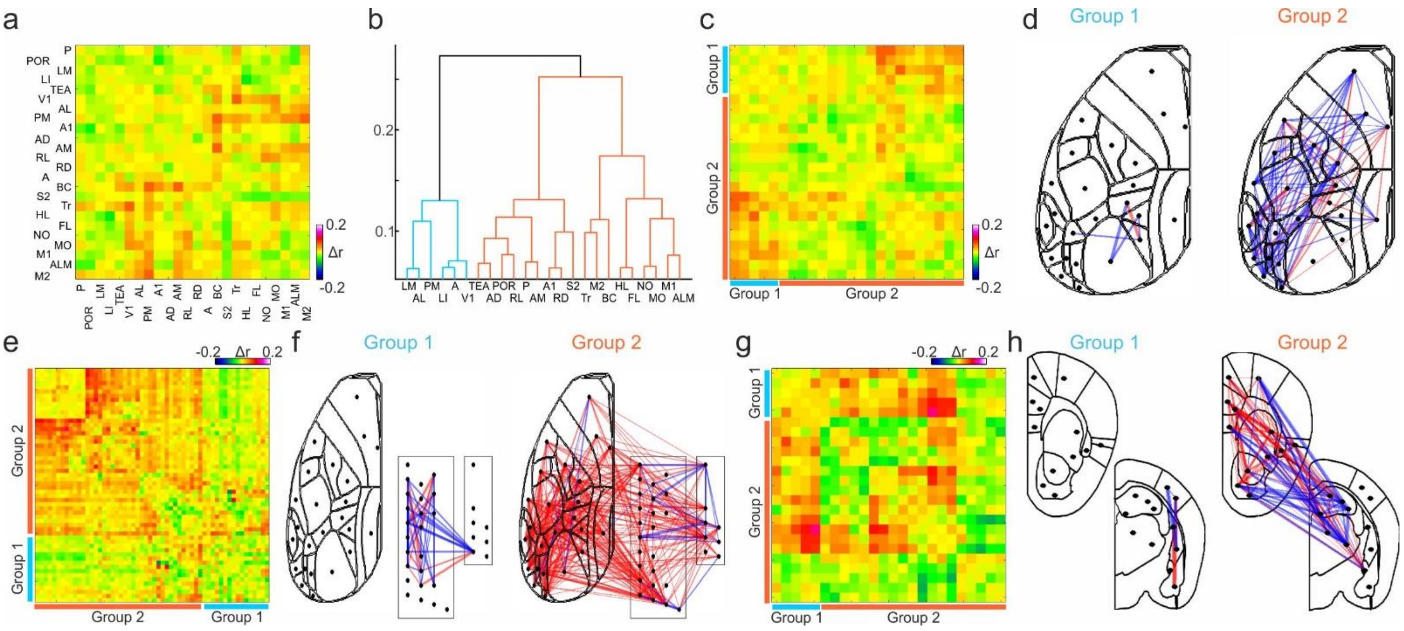


**Figure S3. Correlational subnetworks between 'what' and 'where' tasks. a)** Differential correlation matrix as in Figure 3a, but here we subtracted the 'what' correlation matrix from the 'where' correlation matrix during the sensation period between all cortical areas. Data grouped together from 6 mice. Positive and negative values indicate a bias towards the 'what' and 'where' tasks respectively. **b)** Hierarchical clustering presented as a dendrogram of the differential correlation matrix in a. Two groups are highlighted in different colors. **c)** The differential correlation matrix from a sorted based on the clustering in b. **d)** Pairwise differential correlation values from c superimposed on the cortical atlas for each group separately. Compare to Figure 3d. **e-f)** Similar to c-d, but for the thalamo-cortico-amygdala network. Data grouped together from 4 mice. Compare to Figure 3g. **g-h)** Similar to c-d, but for the subcortical network. Data grouped together from 9 mice. Compare to Figure 3j.

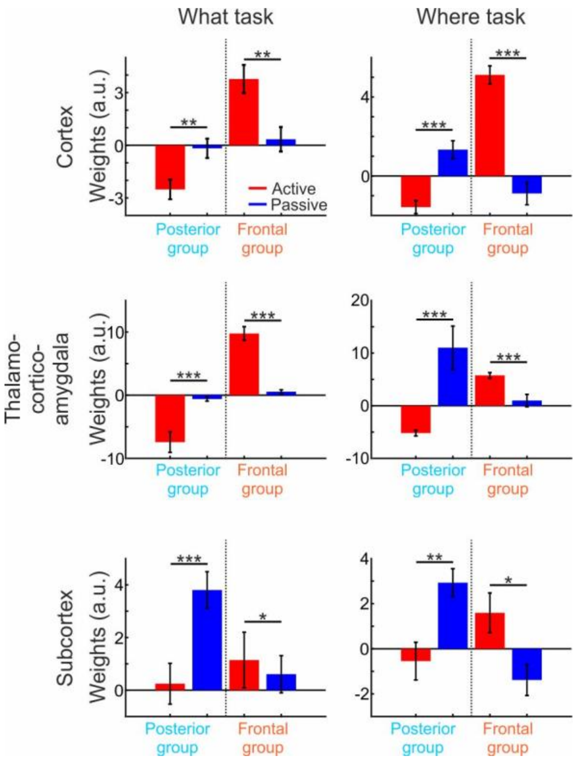


**Figure S4. SVM weights for choice in 'what' and 'where' tasks separately.** Mean weights assigned by a choice SVM (i.e., Hit vs CR; averaged within the sensation period) for active (red) and passive (blue) in the posterior or frontal groups for 'what' (left) and 'where' (right) tasks in cortex, thalamo-cortico-amygdala or subcortical networks (compare to Fig. 3e, g, i). In general, both tasks display similar weight differences. Error bars are SEM over mice (10 iterations from each mouse; n=6 mice for cortex, n=4 for thalamo-cortico-amygdala, n=9 for subcortex). *** - $p<0.05$, *** - $p<0.01$, *** - $p<0.001$ ranked sum test.

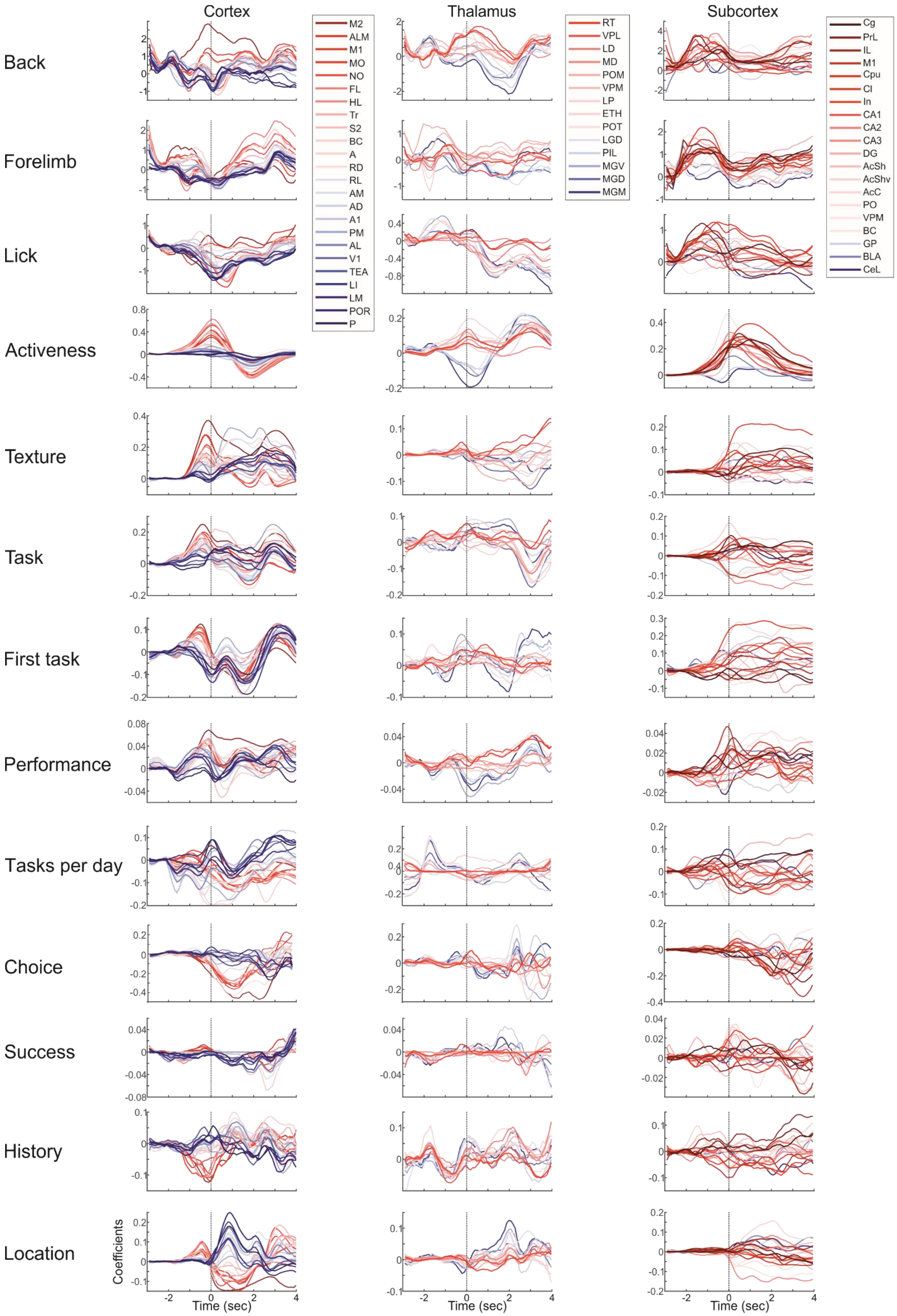
Cortex
Thalamus
Subcortex
Back
Forelimb
Lick
Activeness
Texture
Task
First task
Performance
Tasks per day
Choice
Success
History
Location
Coefficients
Time (sec)
M2
ALM
M1
MO
NO
FL
HL
Tr
S2
BC
A
RD
RL
AM
AD
A1
PM
AL
V1
TEA
LI
LM
POR
P
RT
VPL
LD
MD
POM
VPM
LP
ETH
POT
LGD
PIL
MGV
MGD
MGM
Cg
PrL
IL
M1
Cpu
Cl
In
CA1
CA2
CA3
DG
AcSh
AcShv
AcC
PO
VPM
BC
GP
BLA
CeL

**Figure S5.** Coefficients for each predictor as a function of time derived from the multivariate linear model in Figure 5 in cortex, thalamo-cortico-amygdala and subcortical networks. Red colors indicate frontal areas and blue colors indicate posterior areas.

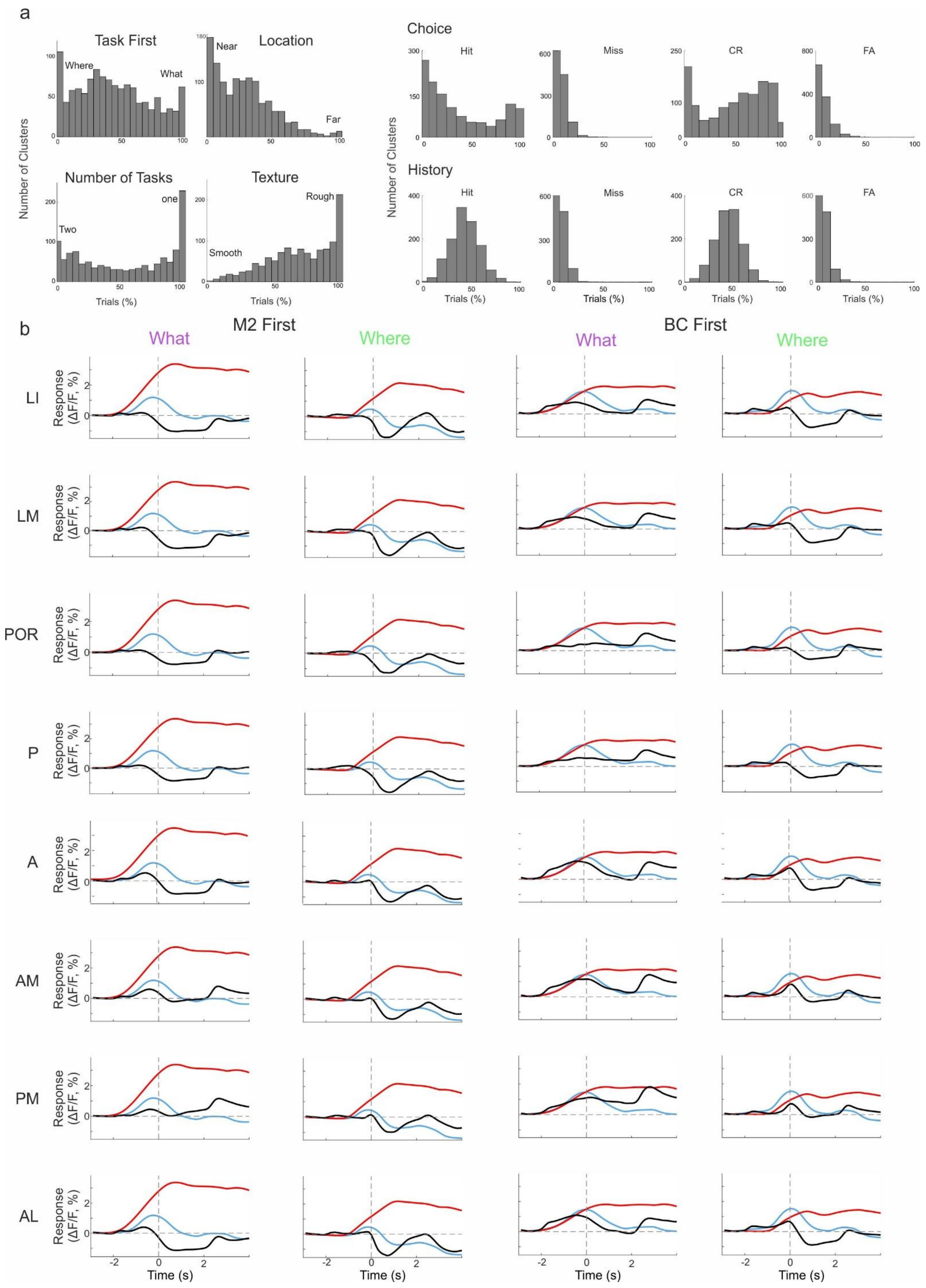


**Figure S6. Autoencoder analysis for different behavioral parameters and responses in different cortical areas. a)** Distribution for the number of clusters as a function of the percentage of trials within a cluster for different behavioral variables (task type and activeness are presented in Figure 6d). **b)** responses in M2 (red), BC (blue) and a specific association area for each of the four groups: M2 first what, M2 first where, BC first what, BC first what.
